# Benchmarking software tools for trimming adapters and merging next-generation sequencing data for ancient DNA

**DOI:** 10.1101/2023.07.17.549303

**Authors:** Annette Lien, Leonardo Pestana Legori, Louis Kraft, Peter Wad Sackett, Gabriel Renaud

## Abstract

Ancient DNA is highly degraded, resulting in very short sequences. Reads generated with modern high-throughput sequencing machines are generally longer than ancient DNA molecules, therefore the reads often contain some portion of the sequencing adaptors. It is crucial to remove those adaptors, as they can interfere with downstream analysis. Furthermore, overlapping portions when DNA has been read forward and backward (paired-end) can be merged to correct sequencing errors and improve read quality. Several tools have been developed for adapter trimming and read merging, however, no one has attempted to evaluate their accuracy and evaluate their potential impact on downstream analyses. Through the simulation of sequencing data, seven commonly used tools were analyzed in their ability to reconstruct ancient DNA sequences through read merging. The analyzed tools exhibit notable differences in their abilities to correct sequence errors and identify the correct read overlap, but the most substantial difference is observed in their ability to calculate quality scores for merged bases. Selecting the most appropriate tool for a given project depends on several factors, although some tools such as fastp have some shortcomings, whereas others like leeHom outperform the other tools in most aspects. While the choice of tool did not result in a measurable difference when analyzing population genetics using principal component analysis, it is important to note that downstream analyses that rely on quality scores can be significantly impacted by the choice of tool.

## 1 Introduction

Next-generation sequencing (NGS) has ushered in a new era of genomics by enabling researchers to sequence DNA at an unprecedented rate, leading to a significant increase in the number of available genome references. Illumina is currently the NGS platform with the highest market share. Library preparation for Illumina sequencing involves three main steps: DNA fragmentation, the addition of adaptors to bind to the flowcell (i.e. the solid medium to allow sequencing), and the amplification of DNA templates for sequencing. DNA templates are then sequenced through the synthesis of complementary DNA strands and optical base calling. Reversible terminator nucleotides are used, with one nucleotide incorporated at a time while capturing its fluorescent signal through high-precision optical imaging. This process is repeated for a specific number of cycles, resulting in read lengths typically ranging from 75 to 250 base pairs (bp). An additional round of sequencing can be performed on the same molecules where the reads are reverse complemented, bound to the flowcell and the nucleotides are read from the other end and the other strand. Reads produced that way are called paired-end reads, the first read is called the *forward* read whereas the second is the *reverse* read. The basecaller is the software used to call nucleotides from raw intensity values. However, due to certain factors, nucleotides can be misread and the basecaller therefore also provides quality scores for each base call, indicating the probability of a sequencing error for that base Liu et al. (2012); Mardis (2008).

One instance where the fragmentation step is not required is working with DNA extracted from fossils, sediments, or museum samples, otherwise called ancient DNA (aDNA). This DNA is naturally degraded and therefore extracted fragments can be on average very short (*<*50 bp). While other idiosyncrasies of aDNA including contamination and chemical damage create specific challenges for downstream analyses, degradation of the DNA molecule needs to be addressed in the earliest stages of the bioinformatics processing Orlando et al. (2021).

While removing lingering sequencing adapters from sequencing reads is important for modern DNA as it can lead to mismappings or misassemblies, it is a crucial step for aDNA. The short fragment size entails that the vast majority of templates will be shorter than the read length and therefore have lingering sequencing adapters, often for the majority of the read (see Figure 1). Furthermore, if the reads are paired-end, after performing adapter trimming, both the remaining forward and reverse reads should be the same length and the reverse complement of each other as it is the same molecule. An additional bioinformatics step is to merge both the remaining forward and reverse reads into a single sequence, as this can correct potential sequencing errors and lead to more accurate data. If paired-end reads are produced and if the length of the template is larger than the read length but still shorter than twice the read length, then both reads should partially overlap (see Figure 1). Similarly, both reads can be merged into a single one, where corrections can be applied to the overlapping portion to mitigate sequencing errors. For aDNA, this bioinformatics step can be construed as reconstructing the original aDNA fragment from either single-end or paired-end reads.

**Figure 1:**
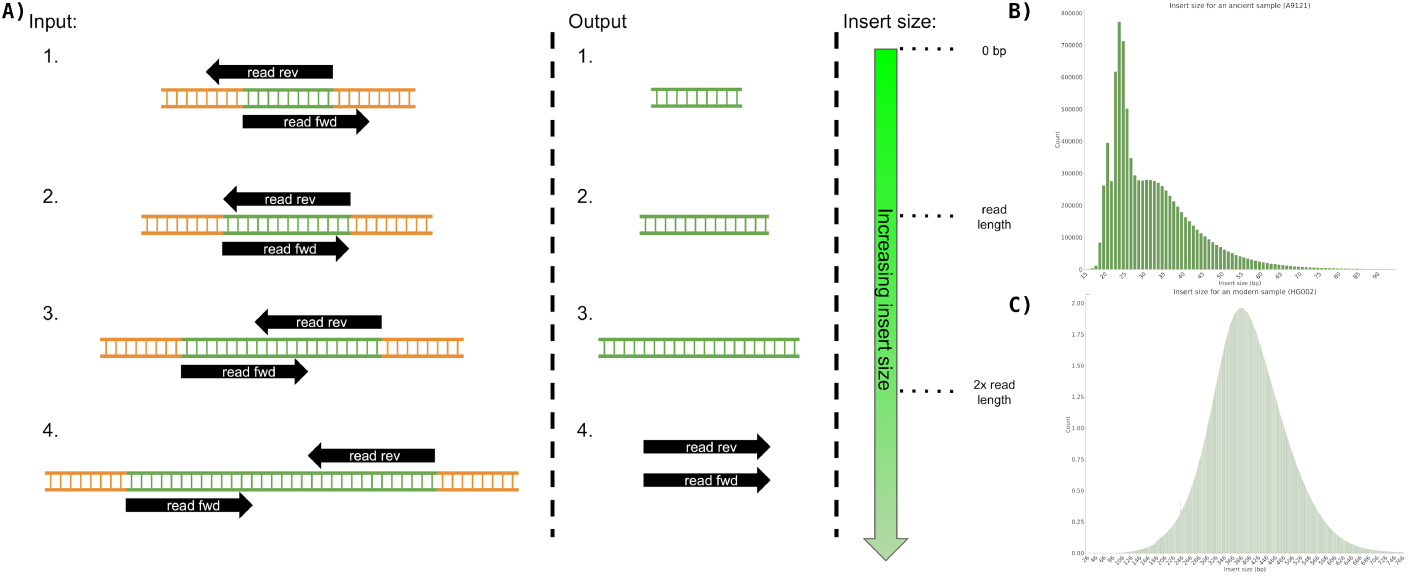
**A)** Example of 4 cases of adapter removal for paired-end sequencing data where the fragment size is: 1. shorter than the read length 2. exactly the read length 3. greater than the read length but shorter than twice the read length, thus resulting in partial overlap 4. more than twice the read length. The desired output for all subcases involves producing a single DNA sequence, except for case 4. where the pairs should be left as they are. B) The fragment size distribution for ancient DNA, data from the A9121 sample from Hajdinjak et al. (2018). C) The fragment size distribution for modern DNA, data from the HG002 sample from Zook et al. (2016)

Several tools have been developed to reconstruct aDNA fragments from sequencing data. These include AdapterRemoval Schubert et al. (2016), bbmerge Bushnell et al. (2017), ClipAndMerge Peltzer et al. (2016), fastp Chen et al. (2018), leeHom Renaud et al. (2014), SeqPrep John (2016), and adna-trim Li (2018). The aforementioned programs have been used in several studies. They use different algorithms, handle mismatches between both reads differently and produce different quality scores. In aDNA studies, often previously published samples from different groups are merged together to answer biologically relevant questions. It is unknown however if different adaptor trimming and read merging strategies lead to batch effects.

In this study, we benchmark different tools which have been designed and used in aDNA studies to reconstruct aDNA from sequencing data. More specifically, using simulated NGS data, we evaluated how these tools perform at various simulated lengths of the template, how robust they are to sequencing errors, and how they compute consensus bases in the presence of matches/mismatches between 2 nucleotides. Finally, we sought to evaluate if using different trimming programs had any impact on the sample placement within a principal component analysis (PCA), which is a very common analysis in aDNA studies.

We find that the analyzed tools exhibit notable differences in their abilities to correct sequence errors and identify the correct read overlap, but the most substantial difference is observed in their ability to calculate quality scores for merged bases. In this regard, we find that fastp might not be suitable for aDNA, whereas leeHom outperforms the other tools in most aspects except for runtime. AdapterRemoval offers favorable accuracy, but has a high rate of false positives. ClipAndMerge and SeqPrep seem to have built-in cutoffs that limit their ability to accurately infer longer templates and lower robustness to sequencing errors. bbmerge seems sensitive to sequencing errors, and its predicted quality scores suffer from poor correlation to their actual error rates. On the other hand, adna-trim offers good accuracy and the best speed but seems to overestimate the error rate of merged portions. While the choice of tool did not result in a measurable difference when analyzing population genetics using principal component analysis, it is important to note that other downstream analyses that rely heavily on quality scores could be significantly impacted by the choice of tool.

## 2 Materials and methods

### 2.1 Generating simulated data

For a detailed description of the generation of simulated data, see Supplementary Information. In brief, we generated simulated aDNA data using the telomere-to-telomere assembly of a human genome T2T-CHM13 and raw sequencing reads for two individuals from the National Institute of Standards and Technology’s Genome in a Bottle (GIAB) project Nurk et al. (2022); Zook et al. (2016). We simulated DNA fragments and paired-end reads using gargammel Renaud et al. (2017) and ART Huang et al. (2012), with varying sequence length distributions, error rates, and Phred quality scores.

### 2.2 Trimming and merging

We compared seven different tools for adapter clipping and paired-end read merging: AdapterRemoval Schubert et al. (2016), bbmerge Bushnell et al. (2017), ClipAndMerge Peltzer et al. (2016), fastp Chen et al. (2018), leeHom Renaud et al. (2014), SeqPrep (John (2016)), and seqtk/adna-trim (Li (2018) and Li (2019)). Only leeHom and seqtk/adna-trim were specifically designed for aDNA analysis, although the merging algorithm of AdapterRemoval is also specifically claimed to be suited for aDNA processing. To enable a representative comparison, we standardized some parameters across the tools. Specifically, the minimum number of overlapping bases was set to 10 and length filtering was disabled across all tools except leeHom, which instead uses a specific parameter for aDNA processing. Detailed descriptions of the parameters used for each tool and results with default minimum overlap and length filtering settings can be found in the Supplementary Information.

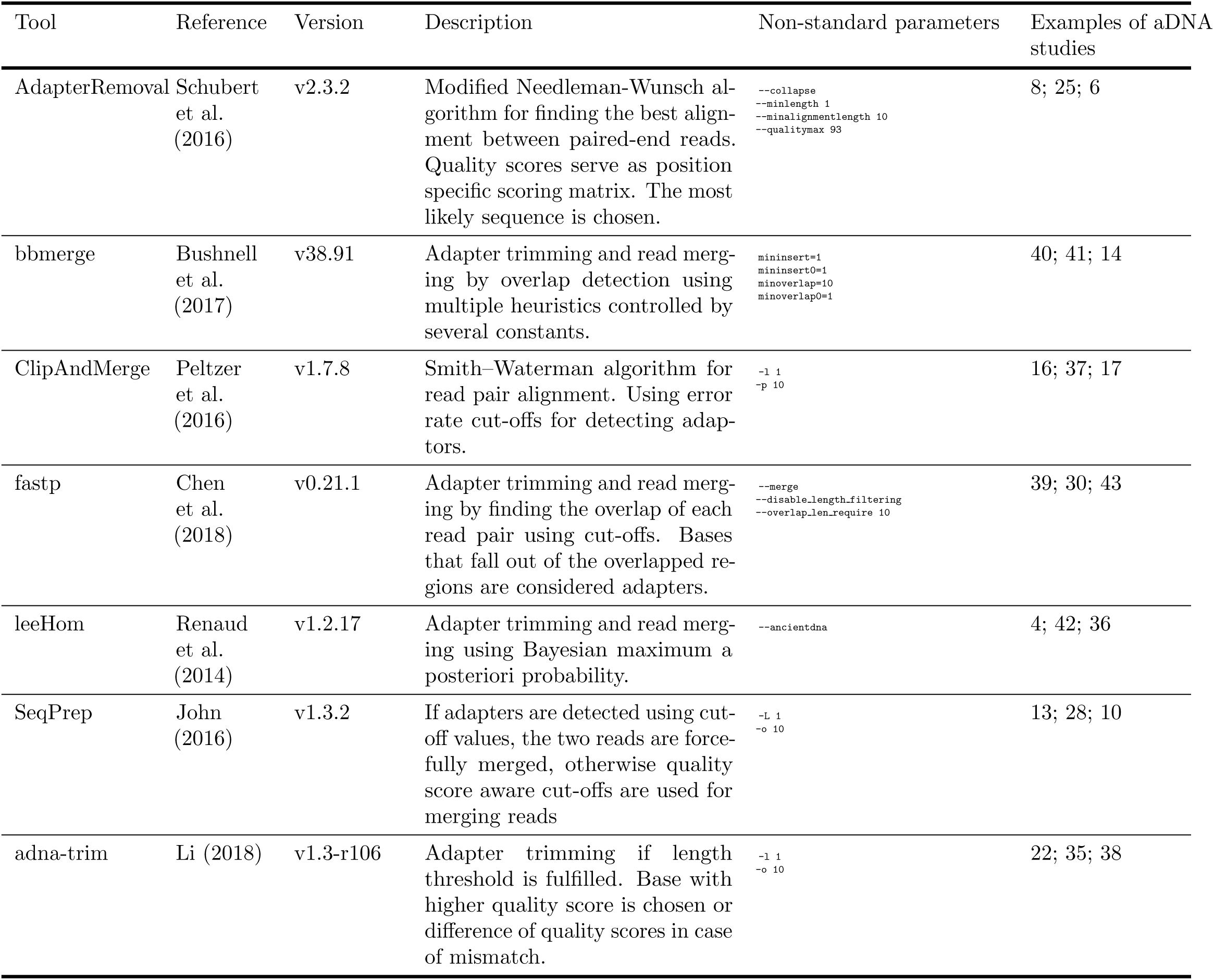

### 2.3 Evaluating base quality score accuracies

We evaluated the accuracies of the merged per-base Phred quality scores by comparing the Phred values to the actual error rates. To calculate the confidence intervals for the observed error rates, we used the Clopper-Pearson method Clopper and Pearson (1934). We then used the inverted ranges of the Phredscaled confidence intervals as weights to calculate a weighted coefficient of determination (*R*^2^) for each tool. For a detailed description of this process, see Supplementary Information.

### 2.4 Population analysis

We compared our aDNA data to the genetic profiles of Eurasian populations from the Affymetrix Human Origins array Lazaridis et al. (2014). Genotyping was performed using BWA mem Li and Durbin (2010), followed by subsampling and pseudohaploid calling with samtools Danecek et al. (2021) and PileupCaller Schiffels (2022). Principal component analysis (PCA) was conducted with smartpca Galinsky and Mah (2022). Within cluster variance was calculated to estimate the influence of the choice of trimming and merging tools on PCA results, more details on its calculation can be found in the Supplementary Information.

### 2.5 Measuring runtime and memory usage

Runtime and memory usage were measured for each trimming and merging tool using snakemake’s Köster and Rahmann (2012) benchmarking functionality. For more details on the machine and memory measurement, please see the Supplementary Information.

### 2.6 Code availability

Code and workflows are available at https://github.com/liannette/mergingtools_ benchmark.

Please refer to the supplement for the complete methodology, including details on reference datasets, genotyping, PCA, within cluster variance calculation, and runtime and memory usage measurement.

## 3 Results

### 3.1 Merge rate and sequence accuracy

Here, a comparative analysis of tools is conducted to evaluate their efficacy in reconstructing DNA insert sequences through paired-end read merging. Differences between the merged read sequence and the DNA insert can arise either from sequencing errors or from incorrect adapter trimming or merging. The measure of sequence dissimilarity is the edit distance, also known as Levenshtein distance, which quantifies the minimum number of single-character edits required to transform one sequence into another. Adapter trimming is generally straightforward, making it a negligible error source, while merged reads with low edit distances are likely to have substitution errors, and those with edit distances larger than 25 may be attributed to incorrect merging. A false negative is defined as a read that should have been merged but was not. Conversely, we have simulated fragment sizes of 1000 bp that should not have been merged, as the reads do not contain any overlap. However, if they were merged, such sequences would be labeled as a false positive.

The accuracy of each tool in reconstructing the sequence of the DNA insert molecules for different insert lengths is depicted in Figure 2.

**Figure 2:**
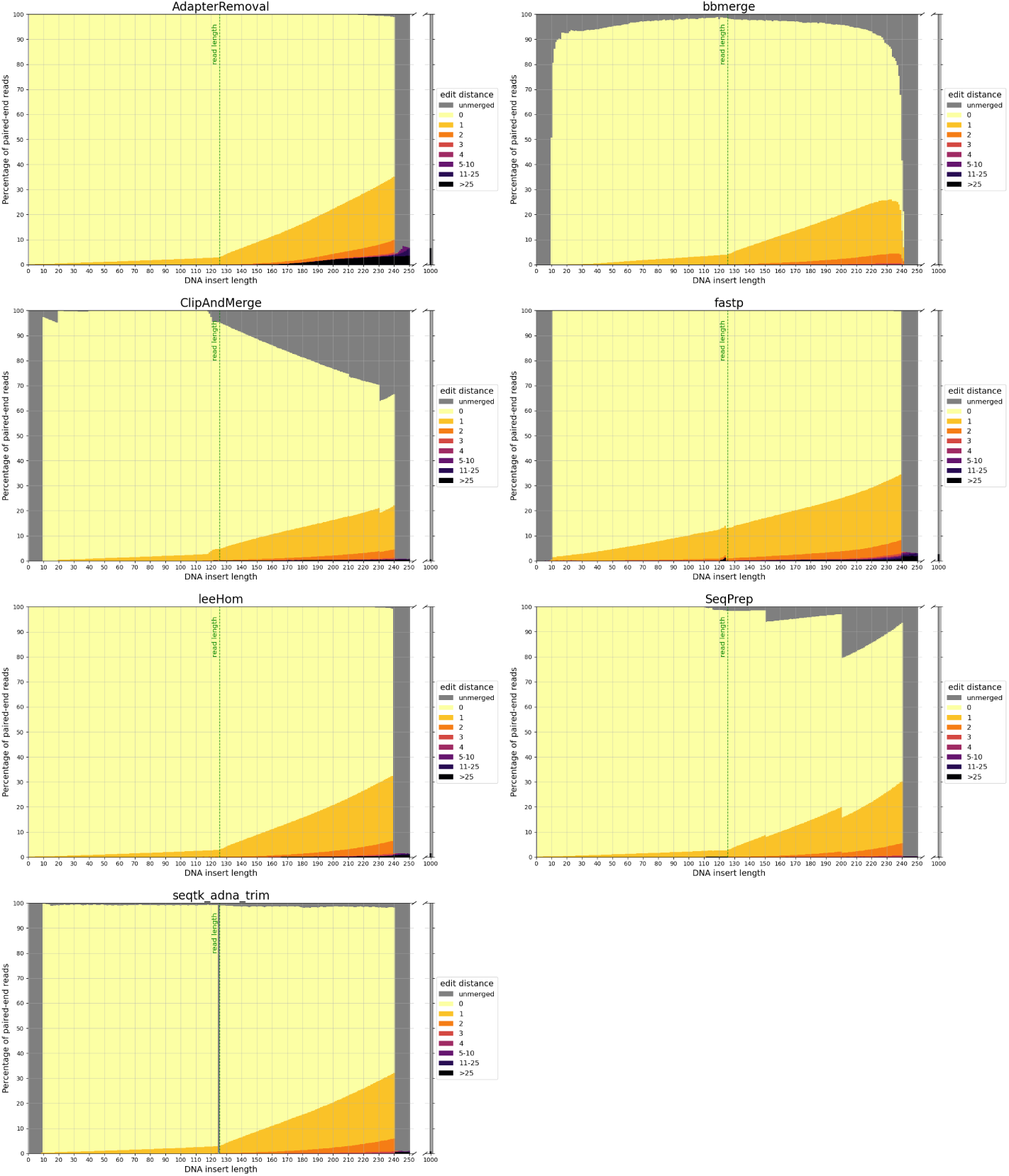
Sequence accuracy of merged reads for different insert sizes. Paired-end reads of DNA inserts with different lengths were simulated and then trimmed and merged with the different tools. High edit distances (in darker colors) correspond to high sequence dissimilarity between the merged read and the DNA insert, and unmerged reads are colored in gray. The green line at 125 bp marks the read length. Paired-end reads of inserts with a length up to this value overlap completely. For higher insert lengths, the read overlap decreases, and the proportion of bases that are unique to only one of the paired-end reads increases. For an insert length of 250 bp and higher, the paired-end reads do not overlap.

#### False-negative merging

To reiterate, paired-end reads that have a sufficient read overlap for merging, but remain unmerged, are classified as false negatives. As the minimum read overlap of merging was standardized to 10 bp (see 2.2), it is not surprising that none of the tools are able to merge reads from inserts with a length exceeding 240 bp. But interestingly, AdapterRemoval, leeHom and SeqPrep are able to merge reads from extremely short inserts that have a length of less than 10 bp, despite them being shorter than the minimum read overlap. Among insert lengths ranging from 10 to 240 bases, high rates of false-negative merging were found for ClipAndMerge and SeqPrep, particularly for insert lengths that surpass the read length. Unmerged reads were also observed across all insert lengths for bbmerge and, to a much lesser degree, for seqtk/adna-trim. For bbmerge, the merge rate is highest when the overlap is maximized, which is the case for inserts the size of the read length. Additionally, seqtk/adna-trim was incapable of merging reads from inserts with a length of 125 bp, equivalent to the read length, which might be due to a bug in the software. False-negative merging behavior was almost non-existent in AdapterRemoval, fastp, and leeHom.

#### False positive merging

In the event that non-overlapping paired-end reads are merged, it is deemed a false-positive. With a read length of 125 bp, the paired-end reads that originate from DNA inserts of 250 bp or more lack any overlapping bases. False-positive merging rates for each tool were calculated as the percentage of merged reads for DNA inserts of 1000 bp length, and are presented both in the gray bars next to all figures in Figure 2 and numerically in Table 1. Among the analyzed tools, AdapterRemoval had the highest false-positive merging rate with over 6 percent, while bbmerge has the lowest false-positive merging rate with less than 0.1 percent. Some tools also incorrectly merge paired-end reads, even when there is adequate overlap for correct merging. Merged reads with an edit distance exceeding 25 for DNA inserts with lengths ranging from 10 to 240 bp are considered to be a result of this. This negative behavior is most pronounced for AdapterRemoval, where the amount of incorrectly merged reads increases with the insert size, ultimately reaching a maximum of over 3 percent for an insert size of 240 bp, indicating that AdapterRemoval is prone to misidentify correct overlap positions. The same can also be observed for ClipAndMerge, fastp and leeHom, but at much lower rates.

**Table 1:** False positive merging rate, calculated as the percentage of merged paired-end reads with no overlap.

| Tool | Estimated false-positive merging rate [%] |
| --- | --- |
| AdapterRemoval | 6.54 |
| bbmerge | 0.0832 |
| ClipAndMerge | 0.597 |
| fastp | 2.65 |
| leeHom | 1.44 |
| SeqPrep | 0.192 |
| seqtk/adna-trim | 0.814 |

### 3.2 Robustness to sequencing errors

This experiment evaluates the ability of each tool in reconstructing the sequences of DNA inserts following different empirical insert length distributions and their robustness to higher rates of sequencing errors. The latter was achieved through quality shift adjustments of the sequencing simulator (further details can be found in the Supplementary Material). The results are illustrated in Figure 3.

**Figure 3:**
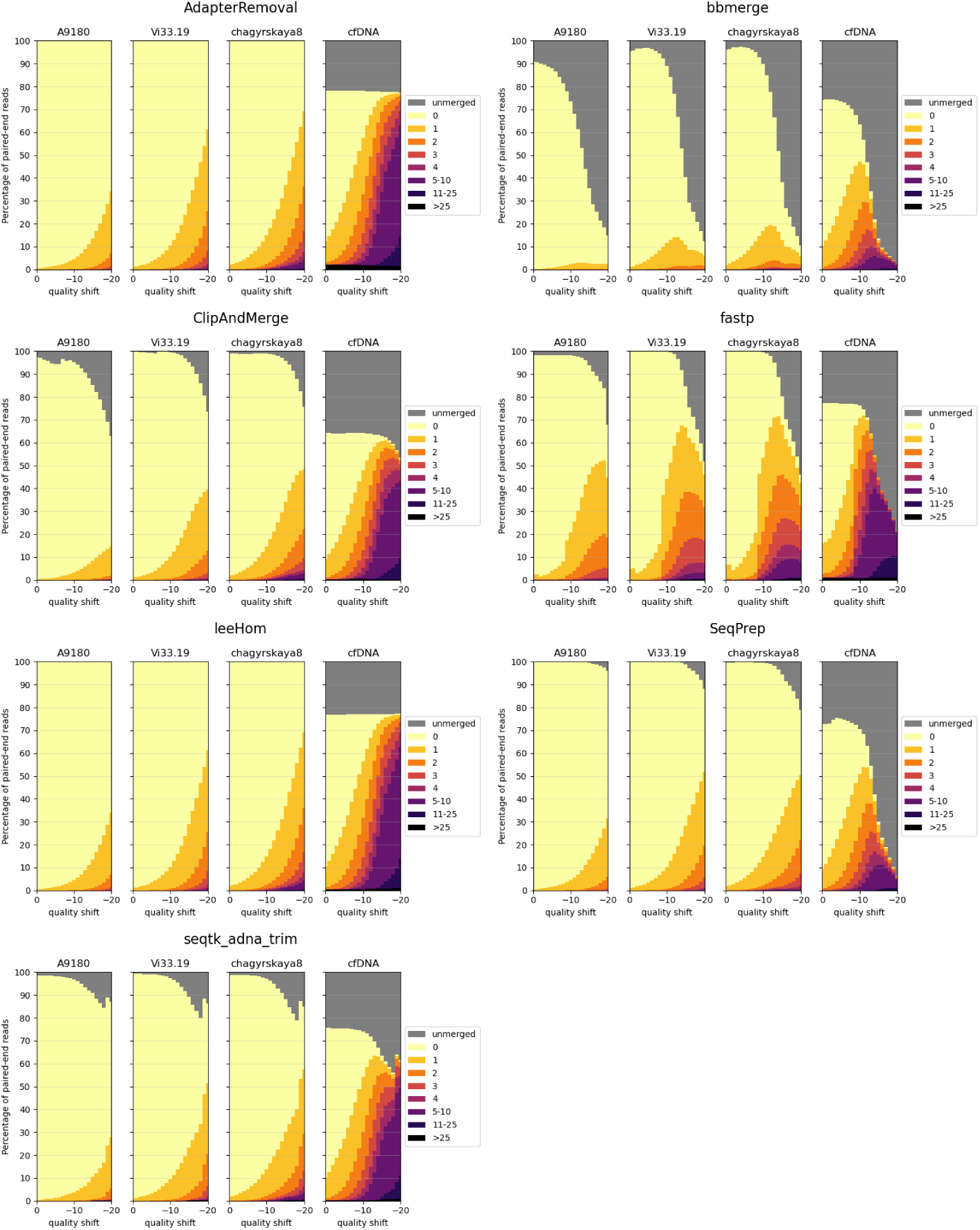
Sequence accuracy of merged reads for different insert size distributions. Paired-end reads of DNA inserts with different length distributions were simulated and then trimmed and merged with the different tools. A9189, Vi33.19 and chagyrskaya8 refer to insert size distributions of aDNA, whereas cfDNA represents the insert size distribution of cell-free DNA. High edit distances (in darker colors) correspond to high sequence dissimilarity between the merged read and the DNA insert.

Three aDNA molecule length distributions (A9180, Vi33.19 and chagyrskaya8) and one distribution of modern, cfDNA were used for this experiment. Histograms of the four insert sizes distributions are depicted in Figure 4. Among the analyzed distributions, A9180 exhibits the shortest insert sizes, predominantly ranging between 10 and 30 bp. It is also the only distribution containing lengths under 30 bp. For Vi33.19 and chagyrskaya8, insert sizes between 40 and 50 bp are most frequent, however, chagyrskaya8 contains also a substantial quantity of lengths exceeding 125 bp, which is not the case for Vi33.19. Most cfDNA molecules span between 125 and 225 bp, with a secondary, smaller distribution at 275-375 bp. At a read length of 125 bp, the vast majority of paired-end reads from aDNA can potentially be merged, whereas for cfDNA, a fraction of reads will remain unmerged due to the lack of overlap.

**Figure 4:**
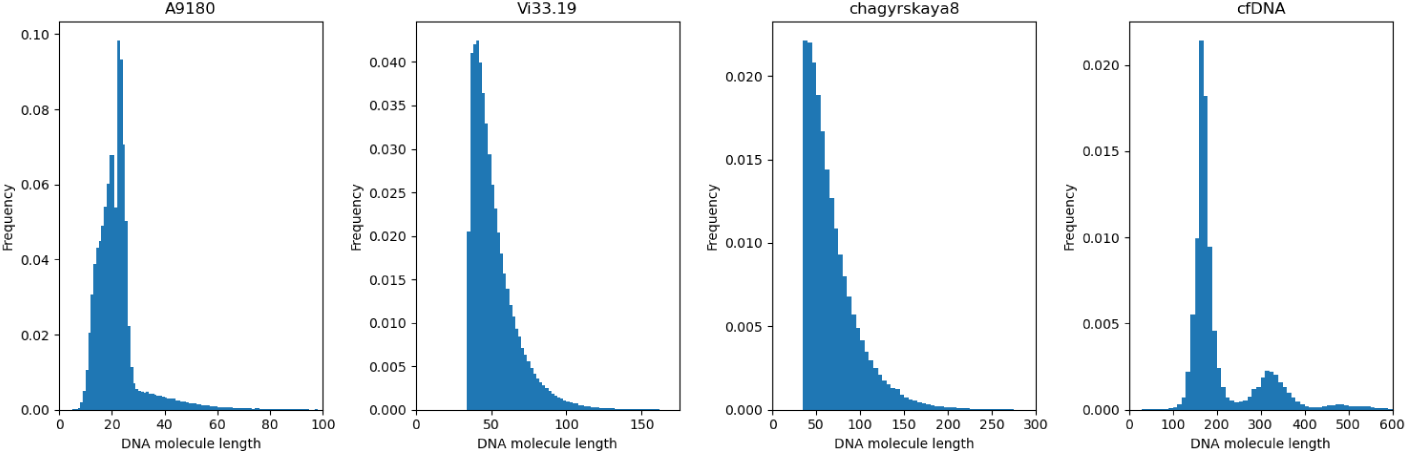
Empirical insert length distributions. A9180, Vi33.19 and chagyrskaya8 are insert size distributions of aDNA generated from different Neanderthal bone samples. cfDNA depicts the insert size distribution of a sample from cell-free DNA.

First, the three aDNA insert size distributions will be examined in detail. At lower sequencing error rates (quality shifts 0 to -10), most tools exhibit similar performance, though some tools fail to merge some reads. This behavior is particularly pronounced for bbmerge. The merged reads with the most errors are generated by fastp. Especially once the sequencing quality falls below a specific threshold (quality shift -9), can an abrupt increase in not-perfectly reconstructed sequences be observed for fastp. AdapterRemoval and leeHom display the highest merge rates and perform almost identically for the aDNA distributions. SeqPrep does not merge some reads at high sequence error rates, but this behavior acts as a sort of filter for reads with a high sequence dissimilarity. In summary, AdapterRemoval and leeHom demonstrate the most robust behavior, meaning that they are capable of merging reads at high sequencing error rates.

Looking at the results for the cfDNA insert size distribution, we see that a significant number of reads remain unmerged across all tools, even at low error rates. This can be attributed to the presence of DNA molecules longer than 250 bp in cfDNA, resulting in paired-end reads with no overlap. ClipAndMerge, however, stands out as it has a higher amount of unmerged reads compared to the other tools. The overall higher amount of longer DNA molecules in cfDNA also accounts for the elevated number of merged reads with high edit distances compared to aDNA, as this not only results in more nucleotides per merged read, but also a smaller read overlap. For bbmerge, fastp and SeqPrep, the merge rate declines dramatically at high sequencing error rates (quality shift of -10 or lower), indicating that these tools are more sensitive to sequencing errors in the overlap. The highest overall merging rate is again exhibited by AdapterRemoval and leeHom. However, of all the tools, AdapterRemoval has the largest number of reads with an edit distance higher than 25 across all quality shifts, indicating that those reads are incorrectly merged.

#### Correcting sequencing errors through merging

The probability of at least one sequencing error occurring within the insert sequence increases with the number of bases in the insert. As a consequence, the percentage of not perfectly reconstructed sequences tends to rise with increasing insert length. Nonetheless, a distinction can be made between paired-end reads with full and partial overlaps. For inserts with a length shorter or equal than the read length, 125 bp, every base pair of the insert is sequenced twice, and as such, is present in both reads. Sequencing errors at any position can, in theory, be corrected by using information from the other read. For inserts that are longer than the read length, some base pairs are, however, only sequenced in one of the reads, making it impossible to correct sequencing errors in those positions. For DNA inserts lengths shorter than the read length, the reads merged with fastp contain significantly more reads with sequencing errors compared to the other tools. For all the analyzed tools, except fastp, the number of reads with edit distances greater than zero increases with the insert length at a slower rate for lengths up to 125 bp, compared to insert lengths exceeding 125 bp.

### 3.3 Per-base merging behavior

In the overlapping region of a paired-end read, each nucleotide base and corresponding Phred quality score is present twice, once on the forward and once on the reverse read. This information is collapsed into a single base and quality score when merging the reads. This experiment evaluates what values each of the tools outputs for any combination of base and quality scores on the forwards and reverse reads.

#### Matching bases

The heatmaps in Figures 5 and 6 visualize the per-base merging behaviors when the base is identical on both the forward and the reverse read. The merging of the base is unambiguous in this case, but the strategies for merging of Phred quality scores vary among the analyzed tools. Some tools generate a new Phred quality score that exceeds either value from the paired-end read, while others select the greater Phred quality score from either read, and some simply select the Phred quality score from the forward read. Sequencing the same base on both reads increases the confidence in the accuracy of that base, thus, the most accurate approach is to increase the Phred quality score in the merged read. Among the analyzed tools, AdapterRemoval, leeHom, and SeqPrep exhibit this behavior. LeeHom and SeqPrep can generate Phred quality scores of up to 60, while AdapterRemoval can even generate values up to 93, although this requires specifying a parameter, as the default maximum Phred value of AdapterRemoval is 41. In contrast, ClipAndMerge and seqtk/adna-trim simply select the greater Phred quality score. The merging behavior of bbmerge lies between the two aforementioned approaches, although it is more similar to the second one. The most simple behavior is displayed by fastp, which just selects the Phred quality score of the forward read and ignores the reverse read entirely.

**Figure 5:**
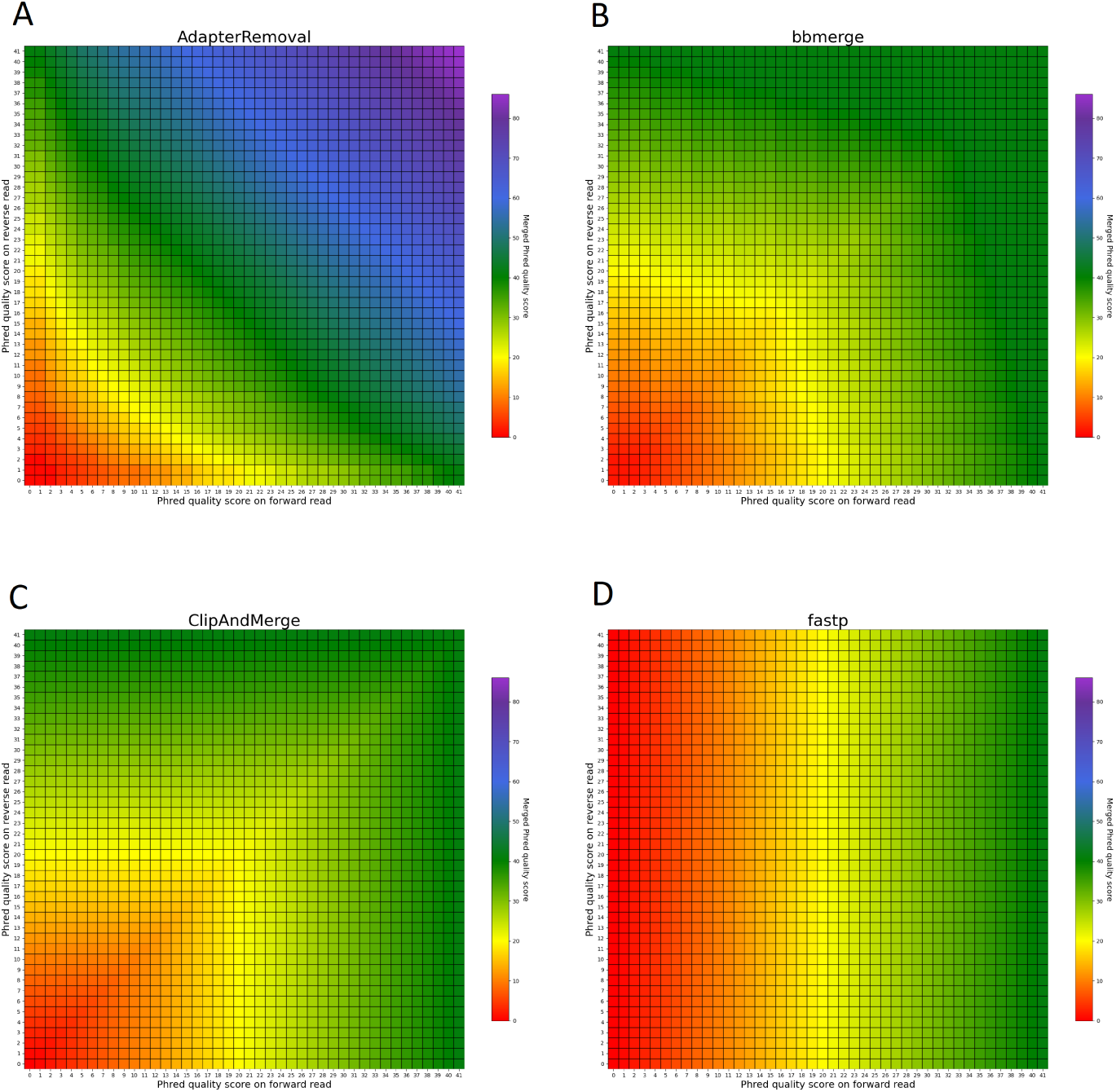
Per-base merging behavior of several tools for matching nucleotides (1 of 2). Results are for (A) AdapterRemoval, (B) bbmerge, (C) ClipAndMerge and (D) fastp. The results for leeHom, SeqPrep, and seqtk/adnatrim can be found in Figure 6. The heatmaps illustrate how distinct Phred quality scores are integrated when overlapping positions with matching nucleotides on both reads are merged. The resulting merged Phred quality score values are color-coded.

**Figure 6:**
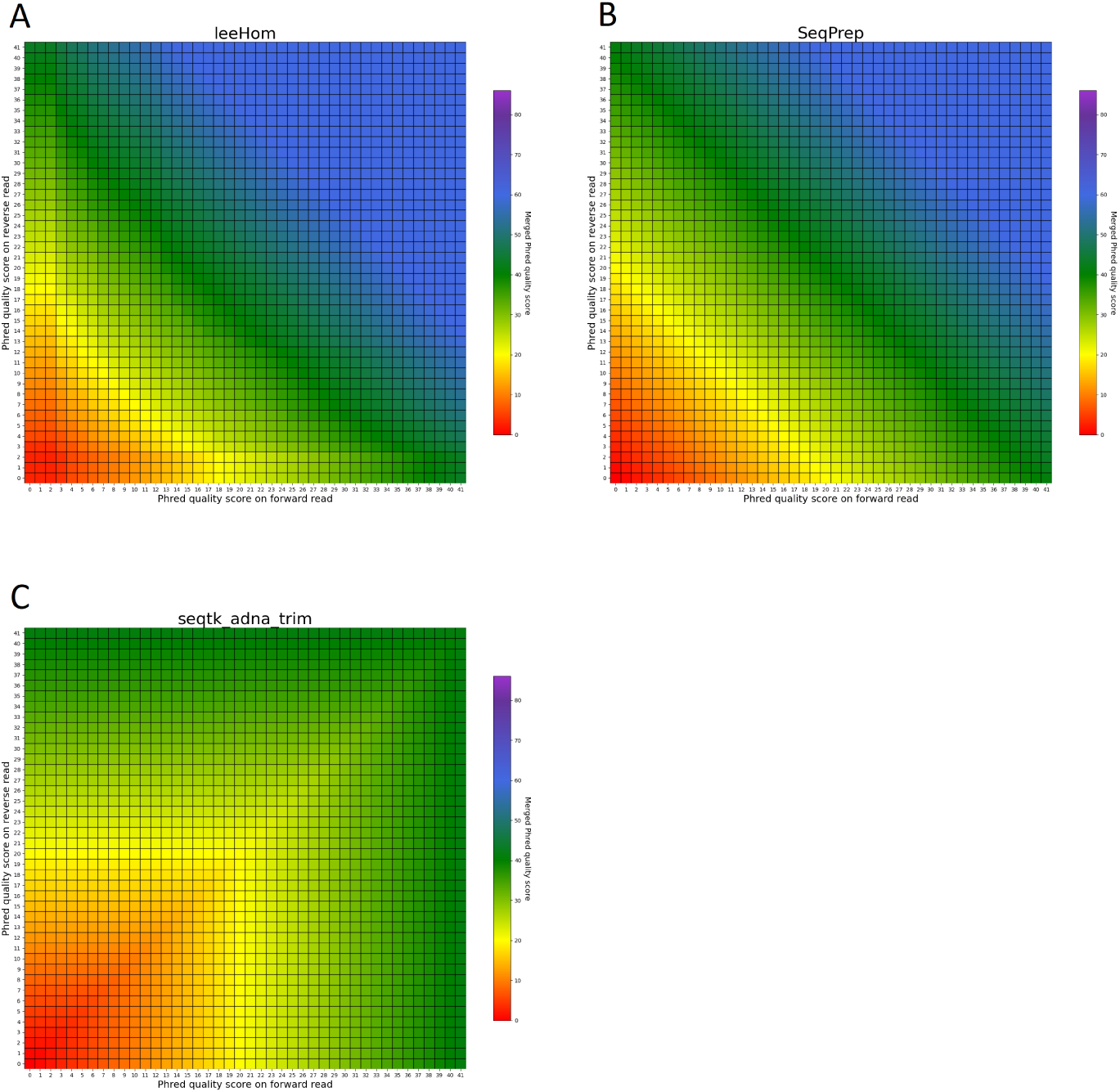
Per-base merging behavior of several tools for matching nucleotides (2 of 2). Results are for (A) AdapterRemoval, (B) bbmerge, (C) ClipAndMerge and (D) fastp. The results for leeHom, SeqPrep, and seqtk/adnatrim can be found in Figure 5. The heatmaps illustrate how distinct Phred quality scores are integrated when overlapping positions with matching nucleotides on both reads are merged. The resulting merged Phred quality score values are color-coded.

#### Mismatching bases

The heatmaps featured in Figure 7 and 8 illustrate the per-base merging behavior of the analyzed tools when the nucleotide base differs between the forward and the reverse read. Again, the tools exhibit distinct behavior patterns in these situations. When a mismatch occurs, the confidence in the accuracy of the merged base should be reduced, particularly when the quality scores of both reads are similar. AdapterRemoval, bbmerge, leeHom, SeqPrep, and seqtk/adnatrim adopt this approach, selecting the base with the highest Phred quality score and using the quality score information from the other read to appropriately adjust the Phred quality score of the merged read. Conversely, ClipAndMerge selects both the base and quality score from the read with the highest Phred quality score, without incorporating information from the other read to modify the Phred quality score in the merged read. fastp, on the other hand, generally obtains the base and quality score from the forward read, except when the quality score is below 15 on the forward read and at least 30 on the reverse read, in which case fastp derives the nucleotide and quality score from the reverse read. This behavior is surprising and indicates that fastp is not as capable as the other tools to reduce sequencing errors.

**Figure 7:**
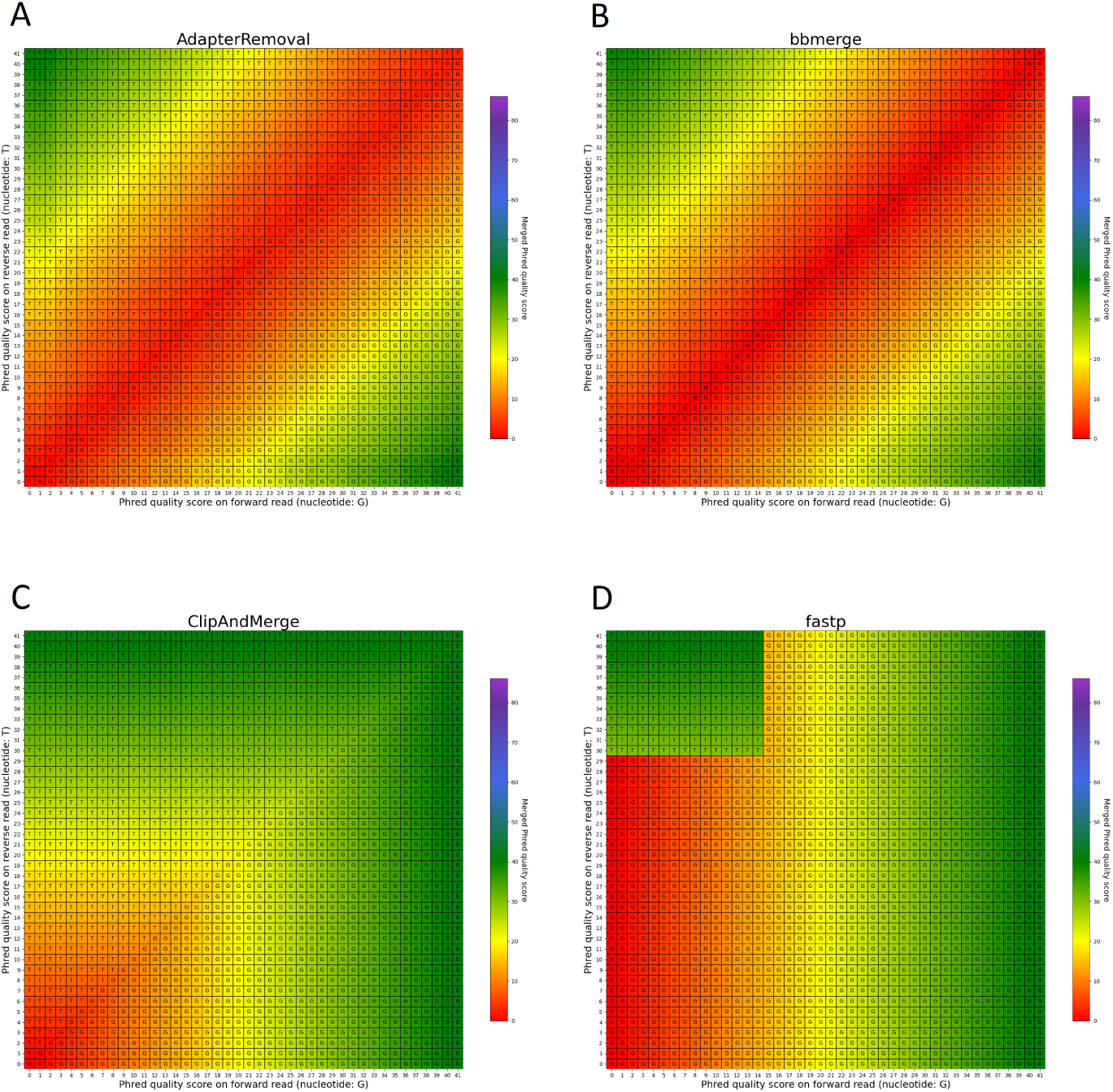
Per-base merging behavior of several tools for mismatching nucleotides (1 of 2). Results are for (A) AdapterRemoval, (B) bbmerge, (C) ClipAndMerge and (D) fastp. The results for leeHom, SeqPrep, and seqtk/adnatrim can be found in Figure 8. The heatmaps illustrate how distinct Phred quality scores are integrated when overlapping positions with matching nucleotides on both reads are merged. The resulting merged Phred quality score values are color-coded, while the letter assigned to each field indicates the nucleotide generated for the merged read. Specifically, when a G is assigned, the base is obtained from the forward read, while a T is derived from the reverse read.

**Figure 8:**
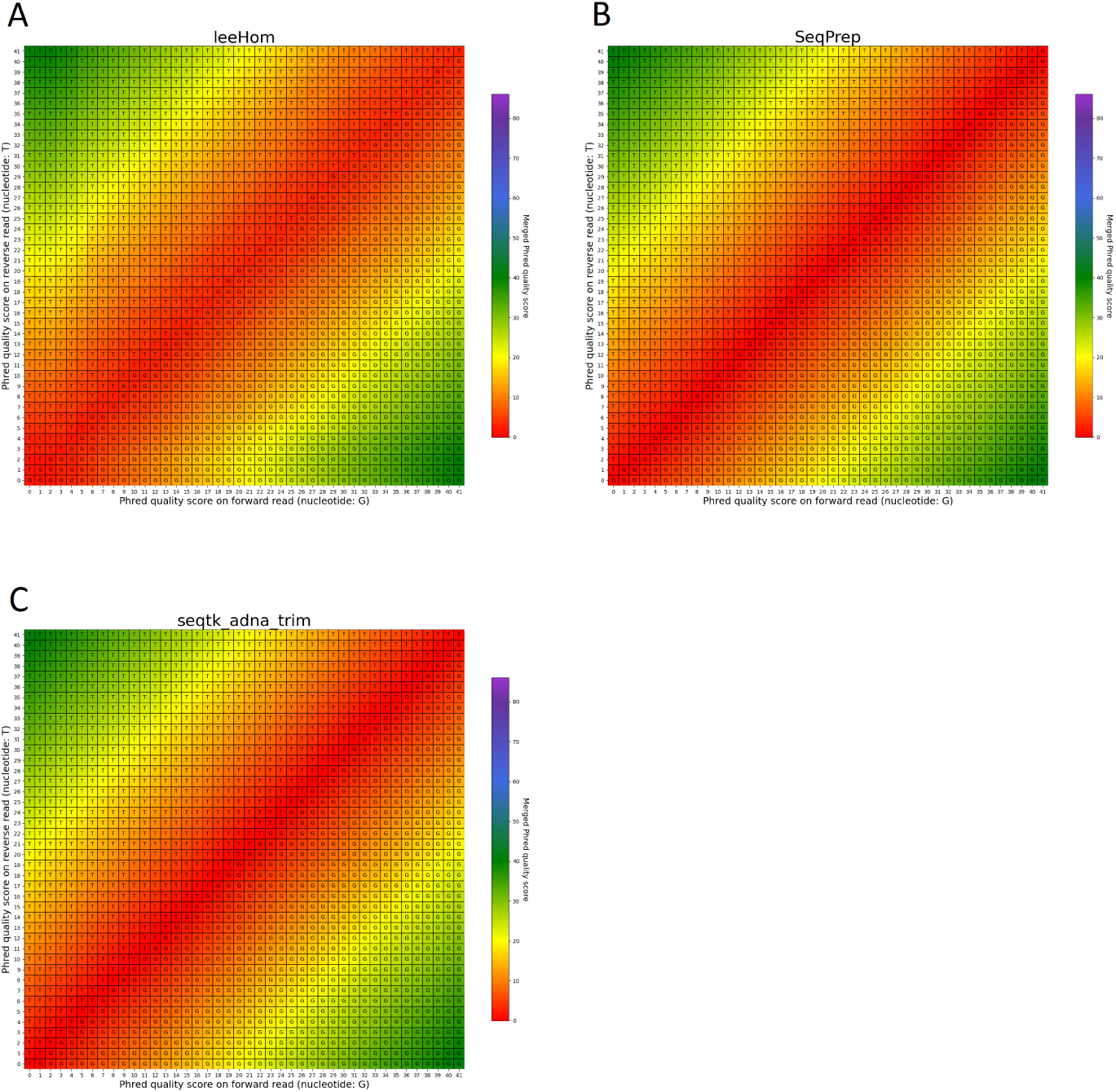
Per-base merging behavior of different tools for mismatching nucleotides (2 of 2). Results are for (A) leeHom, (B) SeqPrep and (C) seqtk/adna-trim. The results for AdapterRemoval, bbmerge, ClipAndMerge and fastp can be found in Figure 7. The heatmaps illustrate how distinct Phred quality scores are integrated when overlapping positions with matching nucleotides on both reads are merged. The resulting merged Phred quality score values are color-coded, while the letter assigned to each field indicates the nucleotide generated for the merged read. Specifically, when a G is assigned, the base is obtained from the forward read, while a T is derived from the reverse read.

In the case of all tools except fastp, an interesting scenario arises when these tools must choose between bases with equal Phred scores during the read merging process. In such instances, AdapterRemoval and leeHom make a random selection between the two bases, while ClipAndMerge and SeqPrep opt for the base from the forward read, and seqtk/adna-trim selects from the reverse read. Notably, bbmerge stands out as the only tool that generates an N in this situation.

### 3.4 Phred quality score accuracy

In this experiment, the Phred quality score accuracy of the tools is analyzed by measuring how close the theoretical error probabilities of the Phred quality scores are to the observable error rates. Figure 9 illustrates the Phred-scaled observed error rates for each merged Phred value, combining the data from five datasets utilizing different quality shifts. Figures displaying the results for each quality shift separately can be found in the Supplementary Information. To indicate how well the Phred values of the observed error rate fit the merged Phred values overall, the weighed coefficient of determination, also called *R*^2^, was calculated for each tool from the combined data, using the inverse range of the confidence intervals as weights to reduce the influence of values with large uncertainty. The weighed *R*^2^ for each of the analyzed tools, whose values can range from 0 to 1, with 1 indicating a perfect fit, can be found in Table 2.

**Figure 9:**
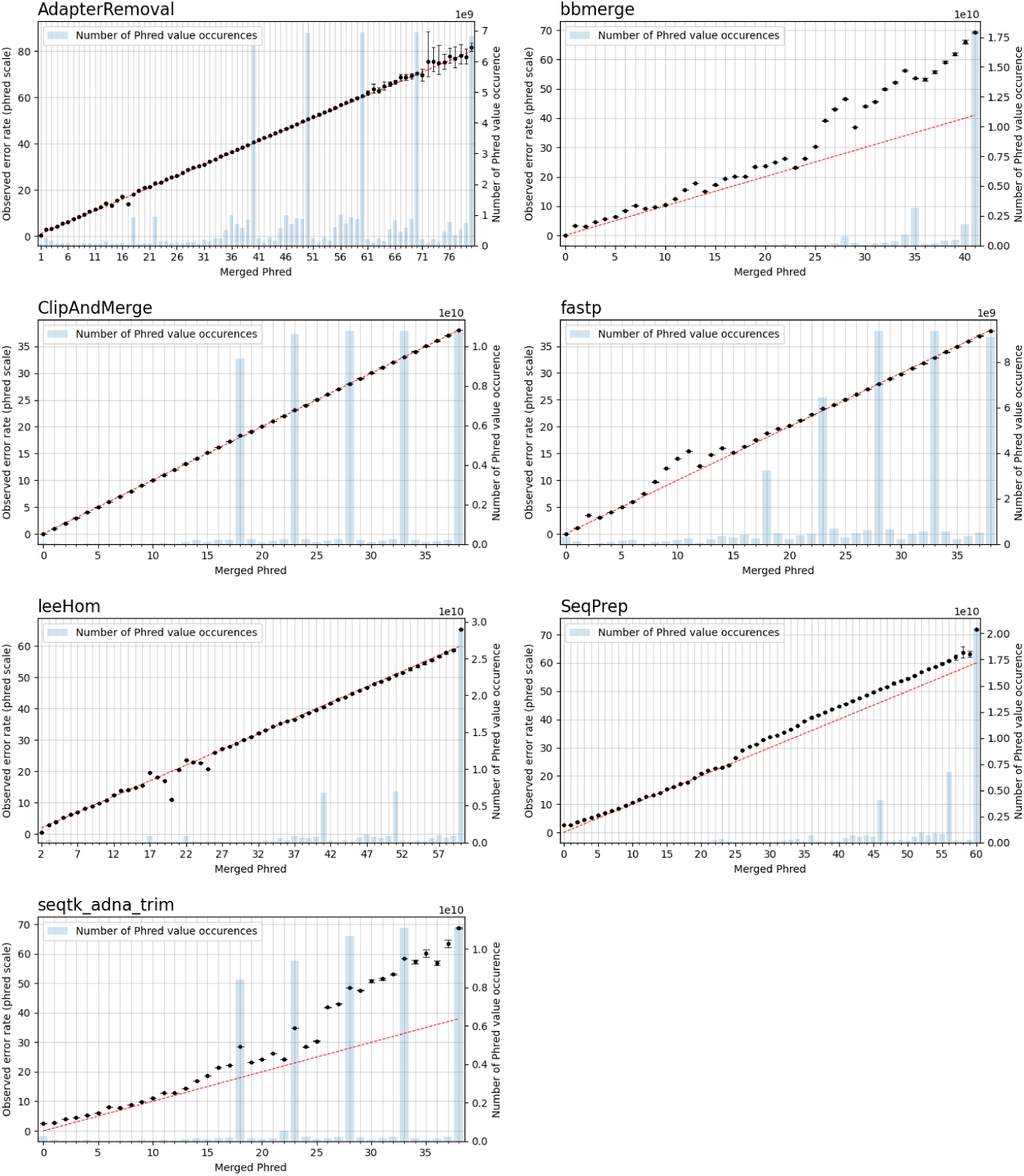
Observable error rates for merged Phred quality score values. The Phred-scaled observable error rates are plotted in black. A high Phred value translates to a low error rate. Confidence intervals for the observable error rates were calculated for *α* = 0.01. The observable error rates were calculated by combining data generated for different simulated sequencing qualities. The red line indicates the perfect fit between observed error rate and merged Phred value, and the number of occurrences of a Phred value in the merged reads is visualized as blue bar plots.

**Table 2:** Weighted *R*^2^ values indicating the error rate accuracy of merged Phred quality scores, calculated by combining data of different sequencing qualities.

| Tool | $R^2$ |
| --- | --- |
| AdapterRemoval | 0.9954 |
| bbmerge | 0.9277 |
| ClipAndMerge | 0.9999 |
| fastp | 0.9943 |
| leeHom | 0.9725 |
| SeqPrep | 0.9707 |
| seqtk/adna-trim | 0.8032 |

The worst fit between the observed error rates and merged Phred values, as indicated by the *R*^2^ value, is seen for seqtk/adna-trim, followed by bbmerge. For both tools, the Phred values of the observed error rates are often higher than the merged Phred values, which means that the actual error rates of merged bases are lower than indicated by their Phred quality scores. For leeHom and SeqPrep, the actual error rate for the (maximum) merged Phred value of 61 is lower than expected. Furthermore, for leeHom and AdapterRemoval more errors were observed for some Phred values around 14 and 25. However, this is only the case at very high sequencing error rates (see Supplementary Results), which is not typically the case with most real data. For fastp, the Phred value calibration is nearly perfect, except for Phred values lower than 15. The best *R*^2^ value is displayed by ClipAndMerge, indicating a near perfect fit of the Phred calibration.

### 3.5 Principal component analysis for population analysis

To examine the potential impact of the different tools, a PCA was performed. Reads of an Ashkenazi Jewish individual (HG002) and a Han Chinese individual (HG005), modified to follow an aDNA insert length distribution, were preprocessed with each of the tools and then genotyped at typical aDNA genome coverages. For each tool and genome coverage, ten samples were generated. Figure 10 shows the first two principal components calculated from the genotype data of present-day Eurasian individuals, on which the HG002 and HG005 samples of approximately 0.25X coverage were projected. All the HG002 and HG005 samples project in the same area as their respective populations, irrespective of the tool used for adapter trimming and read merging. Figures for higher coverages, including zoomed-in plots, in which the samples belonging to the different tools can be distinguished, can be found in the supplement. To estimate the accuracy of each tool when performing a PCA, the variability between the samples was assessed by calculating the within-cluster variance using the coordinates on the first ten principal components. A small within cluster variance indicates that the projections of the samples lie close to each other in the PCA space, which indicates robust results. As seen in Figure 11, the within cluster variance does not significantly differ between the tools, in other words, the choice of tool did not impact the accuracy of the performed PCA.

**Figure 10:**
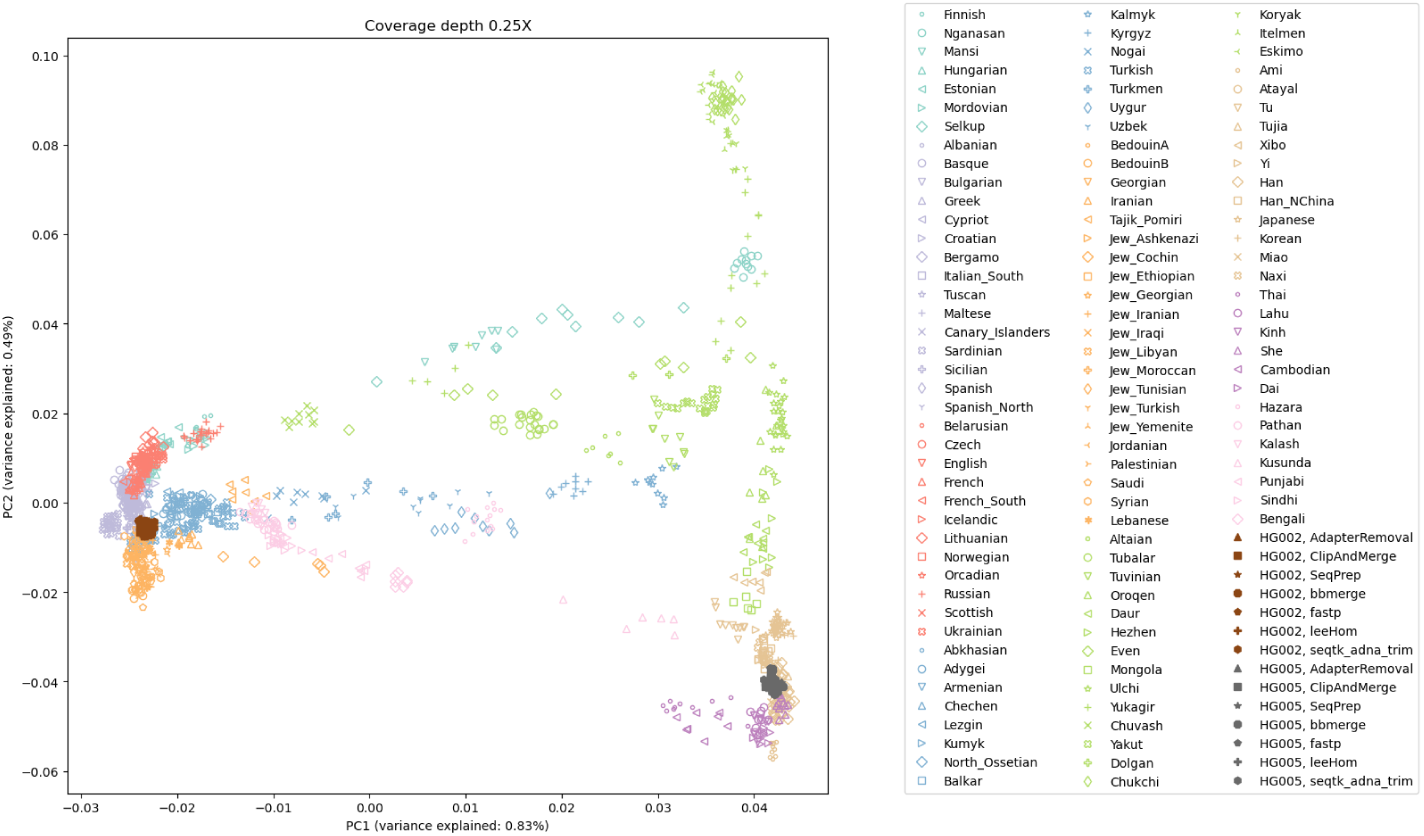
The first two principal components of a PCA of present-day Eurasian individuals; with projected low coverage (0.25X) samples for HG002 and HG005. The projections of all samples for the Ashkenazi Jewish individual (HG002) and the Han Chinese individual (HG005) fall within proximity to their respective populations.

**Figure 11:**
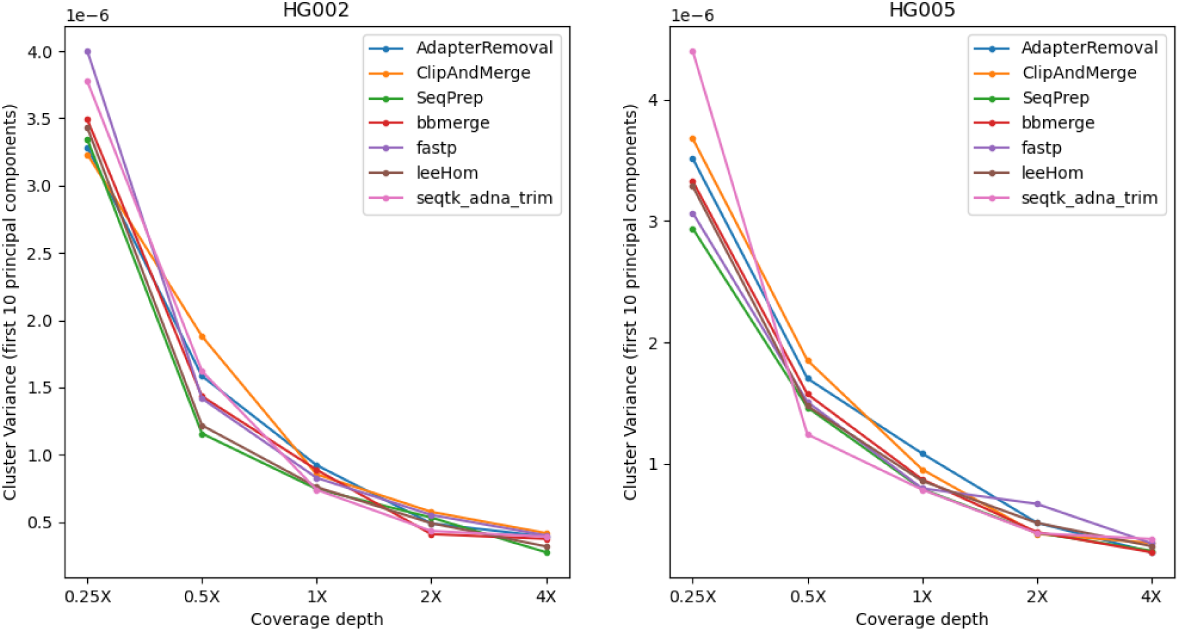
Within-cluster variance of samples projected onto the PCA. Each dot represents the within-cluster variance of 10 data points.

### 3.6 Runtime and memory usage

The runtime and memory usage of each adapter trimming and read merging tool were measured while processing 1 million reads following an aDNA insert size distribution. The tools differ substantially in their resource consumption, shown in Table 3. When comparing the results, it has to be taken into account that ClipAndMerge was utilizing three threads, whereas all other tools were run single-threaded for this experiment.

**Table 3:**
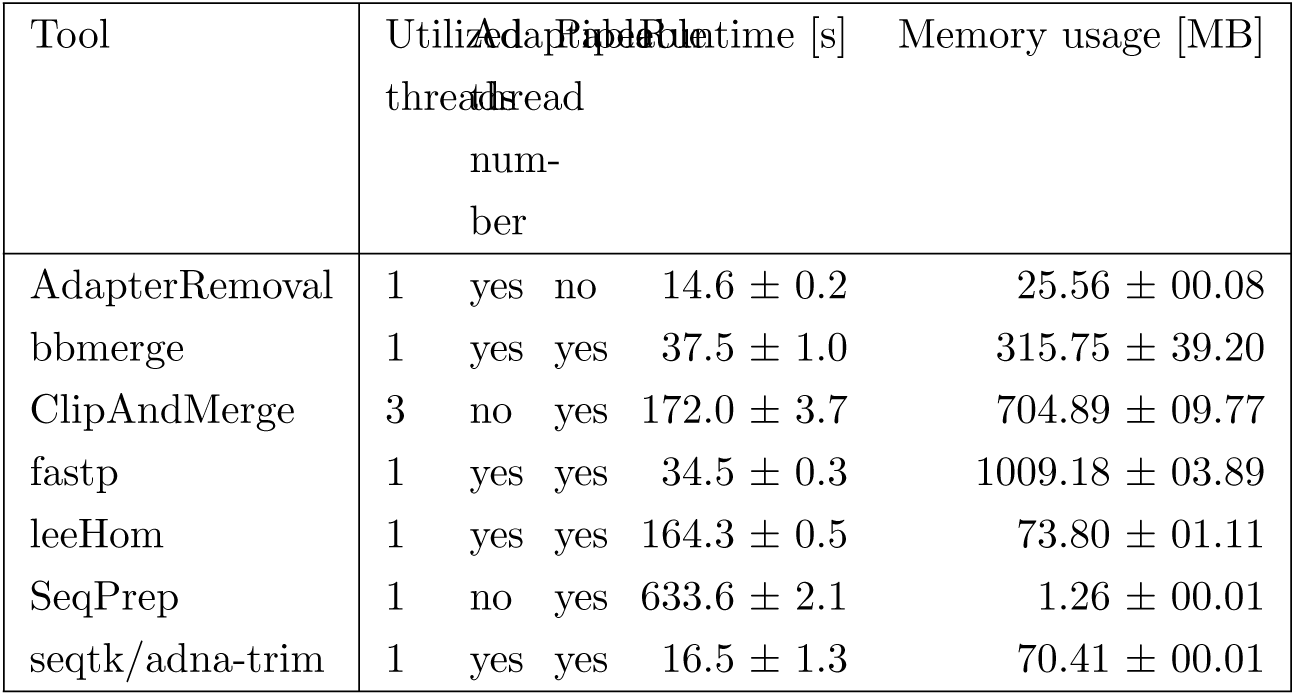
Runtime and memory usage of the analyzed adapter trimming and merging software tools.

Among the tools, fastp stands out as having the highest memory usage among the analyzed tools, followed by ClipAndMerge and bbmerge. The tool with the lowest memory usage is SeqPrep, although it has also by far the longest runtime, almost 4-fold higher than any other tool. A comparably long runtime was also observed for leeHom and ClipAndMerge, despite the latter used three threads. The tool characterized by the shortest runtime is seqtk/adna-trim, followed by fastp, bbmerge and AdapterRemoval. Other considerations related to runtime and memory are the option of piping commands, which not possible with AdapterRemoval, among the possibility of multithreading, which is not supported by SeqPrep, whereas ClipAndMerge number of used threads is fixed at three.

## 4 Discussion

### 4.1 Merged Phred value calculation and accuracy

fastp and ClipAndMerge are considered the least capable among the examined tools in improving read quality, as they always adopt the Phred value from the selected nucleotide instead of calculating a new quality score for a merged base, even when there is a nucleotide mismatch. However, it should be noted that ClipAndMerge generates quality scores that almost perfectly correlate with the actual error rates. It might therefore be relevant for niche applications that depend on highly accurate Phred value calibrations. In contrast, seqtk/adna-trim, and similarly bbmerge, generate a decreased quality score when mismatching nucleotides are merged, but neither of the tools calculates an increased Phred value when merging matching nucleotides. This ambiguous strategy also leads to the least accurate Phred calibrations. The most advanced per-base merging strategies are employed by AdapterRemoval, leeHom, and SeqPrep, as those tools always calculate a new Phred quality score for the merged reads, taking the quality scores from both reads and whether the bases match or not into account. Therefore, those tools should be preferred when the identification of sequencing errors is of importance.

### 4.2 Correcting sequencing errors

Almost all the analyzed tools select the nucleotide with the highest quality score when merging. Only fastp chooses in some cases the base with the lower Phred value, limiting its ability to correct sequencing errors. Specifically, fastp corrects sequencing errors on the forward read only when a high Phred value is found on the reverse read, meaning that sequencing errors on the forward read are not corrected when the sequencing quality is below a certain threshold. Completely overlapping reads merged with fastp contain more sequencing errors compared to the other tools. Furthermore, all tools, except for fastp, exhibit differing rates of sequencing error increase depending on the insert length. This difference is especially apparent when comparing completely overlapping reads to partially overlapping reads, which indicates that some sequencing errors are indeed corrected in the overlapping parts.

### 4.3 Incorrect merging and not-merging

In general, there is a trade-off between incorrectly merged reads and wrongly unmerged reads. For some downstream analyses, incorrectly merged reads would not have a big impact, therefore obtaining the highest possible merge rate is desired. One example would be alignment to a reference genome, where the incorrectly merged reads would either most likely not align at all or result in low alignment scores, making it easy to filter them out. For other applications on the other hand, such as de novo genome assembly, incorrectly merged reads pose a big challenge, and generating fewer reads, but with a higher quality, is preferred.

Among all tools, leeHom and AdapterRemoval have the highest merge rate. Both tools are able to merge almost all reads with sufficient overlap, even at high sequencing error rates. However, AdapterRemoval has the highest tendency to incorrectly merge reads among all tools. Even for paired-end reads that could be merged in theory, AdapterRemoval often misidentifies the correct overlap positions. However, AdapterRemoval’s incorrect merging behavior is primarily present at insert lengths close to double the read length. As aDNA molecules are typically shorter, this aspect has not a big influence when reconstructing the sequences of DNA molecules following the insert size distributions of aDNA. However, the effect is clearly visible when working with DNA insert sizes matching those of cfDNA. A similar effect was observed for ClipAndMerge, but with wrongly unmerged reads (false-negatives) instead of incorrectly merged reads. ClipAndMerge has especially low merge rates for insert sizes between the read length and double the read length (125–250 bp), which is a range that encompasses a significant proportion of cfDNA, but not of aDNA molecules. If the goal is to minimize the number of incorrectly merged reads and sequencing errors, bbmerge would be the best choice. However, bbmerge’s high false-negative merging rate, especially at higher sequencing error rates, makes it a suboptimal choice when working with aDNA, where data is already scarce, or any other analysis where it is desired to merge as many reads as possible. Overall, leeHom most reliably reconstructs DNA sequences, as it is able to merge nearly all reads correctly, even at high rates of sequencing errors. The only caveat is leeHom’s moderate false-positive merging rate for reads with no overlap. Therefore, when working with cfDNA and when presence of incorrectly merged reads is very problematic for further analyses, bbmerge, SeqPrep or seqtk/adna-trim would be a good alternative, however none of those tools are as robust to tolerating sequencing errors in the overlap as leeHom is.

### 4.4 Conclusion

Choosing the appropriate trimming and merging tool for a given project depends on the time and memory that is available, the length of the DNA library molecules, the sequencing quality and what downstream analysis will be performed. In many aspects, leeHom outperforms the other tools. Not only displays leeHom the most statistically correct method in calculating merged Phred values, it also has a high merging rate, mostly identifies the read overlap correctly and is robust for sequencing errors in the overlap. The runtime of leeHom is longer than those of most other tools, but it is able to run as a piped command. While AdapterRemoval is in many ways similar to leeHom, it produces a high amount of incorrectly merged reads. Only in cases where it is preferred to have a reduced amount of merged reads in favor of minimal incorrectly-merged reads, bbmerge would be the tool of choice. Some tools are characterized by especially large resource usage. While for SeqPrep the runtime is exceptionally large, the memory usage of fastp is exceptionally large. fastp also displays inferior sequence correction capabilities and should be avoided. Furthermore, ClipAndMerge’s algorithm to calculate the quality score does not incorporate mismatch information and should also be avoided if the identification of sequencing errors is important for further analysis steps. Although the analyzed tools differ profoundly in their algorithms and the resulting ability to reconstruct aDNA sequences, they had no measurable effect when performing a PCA, which was used to assess influences on downstream analysis. However, other downstream analyses that rely on Phred quality score, should be tested in future experiments, as this is the main aspect in which the tools differ from each other.

## Acknowledgments

This research project and the PhD scholarships of LK were funded by the Novo Nordisk Data Science Investigator grant (number NNF20OC0062491). We would like to thank the Department of Healthtech at The Technical University of Denmark for additional funding.

## Supplemental Data

Supplementary Material can be found online.

## Notes

### Competing Interest Statement

The authors declare that the research was conducted in the absence of any commercial or financial relationships that could be construed as a potential conflict of interest. GR is the programmer behind leeHom. However, AL and LPL did not treat this program any differently from the others.

