## Supplemental Material for "Benchmarking software tools for trimming adapters and merging next-generation sequencing data for ancient DNA"

### 1 Supplementary Methods

#### 1.1 Generating simulated data

##### 1.1.1 Reference sequences

For the first three experiments, DNA sequences were sampled from the telomere-to-telomere assembly of a human genome T2T-CHM13 to simulate human DNA molecules of varying length. The T2T-CHM13 genome was derived from the CHM13 cell line and includes gapless assemblies for all chromosomes except Y, containing 3.055 billion base-pairs in total Nurk et al. (2022).
For the PCA experiment, raw sequencing reads for two individuals were acquired from the National Institute of Standards and Technology's (NIST) Genome in a Bottle (GIAB) project Zook et al. (2016). The first individual, HG002, is a male of Ashkenazi Jewish ancestry, while the second individual, HG005, is a male of Han Chinese ancestry. The DNA was extracted from a large homogenized growth of B lymphoblastoid cell lines from the Personal Genome Project, prepared into a whole genome sequencing (WGS) paired-end PCR-free library and sequenced with an Illumina HiSeq 2500 Rapid SBS system, resulting in 2×250bp reads Zook et al. (2016). Only the forward reads were used for this experiment.

##### 1.1.2 Insert length distributions

For some experiments, sequences matching a specific sequence length distribution of DNA molecules were simulated. Sequencing data from three different ancient DNA (aDNA) libraries, A9180, Vi33.19, and chagyrskaya8, and of one modern cell-free DNA (cfDNA) library were used to obtain the insert size length distributions. The three aDNA distributions contain 1 million insert lengths each, while the cfDNA distribution contains 548767 insert size lengths. The A9180 library Hajdinjak et al. (2018) was generated from an 44,600-43,000 years old skull fragment of an 2-3 year old Neanderthal infant, Mezmaiskaya 2, found in the Mezmaiskaya cave in the Russian Caucasus Pinhasi et al. (2011). The Vi33.19 length distribution belongs to DNA sequenced from a Neanderthal bone sample, which was found in the Vindija Cave in Croatia and was dated to be older than 45,500 years Prüfer et al. (2017). Chagyrskaya8 is a Neanderthal from Chagyrskaya Cave, which is estimated to have lived around 80,000 years ago and the DNA that was extracted from a bone sample Mafessoni et al. (2020). The last insert size distribution, the one for cfDNA, was added, because cfDNA, which refers to all non-encapsulated DNA in the blood stream, also contains a high amount of short DNA molecules. The experiments might therefore also be relevant for analyzing cfDNA, even if it is not the main focus of this analysis.

##### 1.1.3 Simulating DNA fragments and paired-end reads

The adapter sequences used for simulating the paired-end reads were AGATCGGAAGAGCACACGTCTGAACTCCAGTCACCGATTCGATCTCGTATGCCGTCTTCTGCTTG for the forward read and AGATCGGAAGAGCGTCGTGTAGGGAAAGAGTGTAGATCTCGGTGGTCGCCGTATCATTT for the reverse complement read. Except for the experiment regarding per-base merging behavior, sequences of the DNA fragments were simulated with gargammel Renaud et al. (2017), and ART (ART_Illumina (2008-2016), Q Version 2.5.8 from Jun 6, 2016) Huang et al. (2012) was used to simulate the sequencing reads. Sequencing errors and quality scores were introduced according to the profile of the Illumina HiSeq 2500 System (*–seqSys HS25*), with the constriction that no insertion or deletion errors were permitted (*-ir 0*, *-ir2 0*, *-dr 0* and *-dr2 0*). In the cases where higher rates of substitution errors were introduced, the quality shift parameter (*-qs*) was adjusted. Shifting the default value of 0 value by $x$, does not only result in an error rate of $1/10^{x/10}$ of the default error profile Prüfer et al. (2017), it also shifts the Phred values by $x$, as can be seen in figure 1.

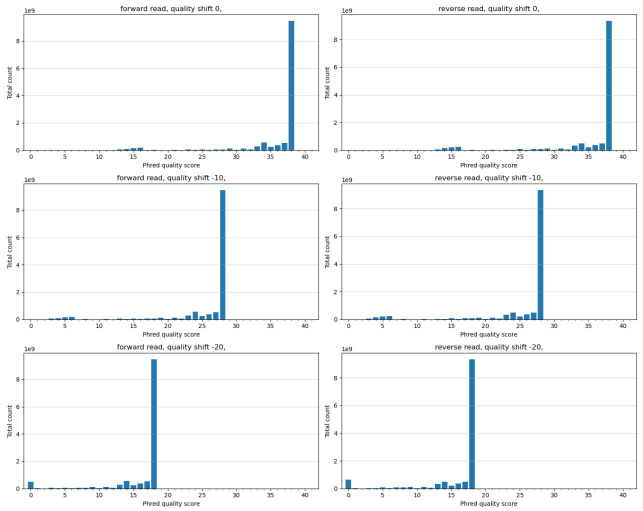


Figure 1: Per-base Phred quality scores distribution of paired-end reads generated with ART with different quality shifts.

**Simulated data generation for per-base merging behavior**
To conduct this experiment, a set of paired-end reads was generated using a custom script. These reads were created by adding adapter sequences to a 32 bp long DNA sequence obtained from the human genome, T2T-CHM13, (chromosome 4, position 155073817:155073849), resulting in a total read length of 75 bp. The reads were designed to differ only at the 16th position of the DNA sequence. The DNA sequence is ACCTCCCAAGCTCAAGTGATCCTCCCACCTC on the forward read, with GAGGTGGGAGGATCACTTGAGCTTGGGAGGT on the reverse read for a match and GAGGTGGGAGGATCA**T**TTGAGCTTGGGAGGT for a mismatch. At variable position, every combination of Phred quality scores ranging from 0 to 41 between the forward and reverse read was presented by one read pair. This was done for both a match and for a mismatch. At all other positions, a Phred quality score of 41 was assigned.

**Simulated data generation for merge rate and sequence accuracy**
For each DNA molecule length between 1 and 250 bp, plus 1000 bp, and chosen insert length distribution (1.1.2), 1 million DNA fragments have been simulated by sampling sequences at random from the T2T-CHM13 human genome using the gargammel subprogram fragsim. In the next step, gargammel's subprogram adptsim was utilized to add adapter sequences, using paired-end mode and a read length of 125 bp. Finally, ART was used to generate paired-end reads with sequencing errors and Phred quality scores. For the reads that were simulated using the insert length distributions, increasing rates of sequencing errors were introduced (quality shift values 0 to -20), in order to test the robustness of the trimming and merging tools to higher sequencing error rates.

**Simulated data generation for phred quality score accuracy**
The data was simulated similarly as for the previous described experiment. To simulate the DNA inserts, 100 million DNA sequences with a length of 120 bp were sampled at random positions from the T2T-CHM13 human genome using the gargammel subprogram fragsim. Then, adapter sequences were added using gargammel's subprogram adptsim, specifing paired-end mode and a read length of 125 bp. Finally, paired-end reads with sequencing errors and Phred quality scores were generated with ART. This was done for different quality shift values (0, -5, -10, -15 and -20), to generate enough data for each Phred quality score value. All the simulated paired-end reads overlap completely, because the simulated DNA inserts are shorter than the read length.

**Simulated data generation for population analysis (PCA)**
To generate simulated paired-end reads for the PCA for the population analysis, raw forwards reads with a length of 250 bp/read of two modern, human males were used as starting material. AdapterRemoval (version 2.3.2) was used to trim any adapters sequences and to filter reads shorter than 250 bp (*–minlength 250*). This removed any reads containing adapter sequences. Using gargammel fragSim, the reads were then trimmed according to the aDNA insert length distribution of Vi33.19. The distribution contained a small fraction (less than

0.001%) of lengths longer than 250. Those lengths were changed to 250, as it was not possible to generate longer sequences from the 250 bp long reads. The reads were then transformed to fasta format with seqkit's subprogram fq2fa Shen et al. (2016), before adapter sequences were added using gargammel's subprogram adptsim (paired-end mode, read length 125 bp) and ART was used to generate the final paired-end reads with simulated sequencing errors and Phred quality scores.

#### 1.2 Trimming and merging

To enable a reasonable comparison, the following parameters have been standardized for all tools except leeHom: the minimum number of overlapping bases for merging was set to 10 and no length filtering was disabled or the minimum length was set to 1. For leeHom a specific setting for aDNA analysis was used. For AdapterRemoval, the parameter for the maximum quality score was set to its maximum value of 93. For adna-trim, an interleaved fastq file was generated with seqtk (version 1.3-r106) using the *mergepe* command. The output was then piped to adna-trim is used for adapter trimming and merging. adna-trim does not take adapter sequences as input, but they were added as input for all other tools.
When analyzing the per-base merging behavior of bbmerge, some filter settings were disabled (*efilter=-1* and *ratiomargin=1*), in order to obtain a complete dataset.

#### 1.3 Evaluating base quality score accuracies

To estimate how well the Phred quality score values for each merged base conform with the error rates they represent, only merged reads that have the right sequence length are considered. This ensures that each position of the merged read can be compared to the same position of the DNA insert. The number of occurrences in the merged reads and the number of times they were associated to a base that did not match the base in the original sequence was counted for each Phred value. From this, the observable error rate is calculated together with a confidence interval, and then translated to the Phred scale. The exact binominal confidence interval was calculated using the method described by Clopper and Pearson (1934), utilizing a two single-tailed Binomial test to calculate the lower and upper bounds of the confidence interval. With this method, confidence intervals can be generated for proportions close to 0, which was the case for most values. The inverted ranges of the Phred-scaled confidence intervals were then in turn used as weights to calculate a weighted coefficient of determination (R-squared) for each tool. This is done to ensure that the values with a large confidence interval have less influence on the R-squared value. The weighted R-squared value indicates how well the error probabilities of the merged Phred quality scores overall correlate with the observed error rates.

#### 1.4 Population analysis

##### 1.4.1 Reference dataset

The data generated for the two individuals was compared against the genetic profiles of Eurasian populations genotyped in the Affymetrix Human Origins array as described in Lazaridis et al. (2014).

##### 1.4.2 Genotyping

Using BWA mem (version 0.7.17-r1188) Li and Durbin (2010), the merged reads were aligned against the 1000genomes Phase2 Reference Genome Sequence (hs37d5), based on NCBI GRCh37, which is compatible with the SNP positions specified in the Affymetrix Human Origins array. PCR duplicates were marked using the MarkDuplicates program of Picard (version 2.27.5) Pic (2019). The alignment was then subsampled using samtools's (version 1.13) Danecek et al. (2021) view command to simulate overall genome coverages of approximately 0.25X, 0.5X, 1X, 2X and 4X. For each coverage, ten subsampled alignment files were generated using different seed parameter values. For the SNPs positions that are present in the Affymetrix Human Origins panel, a pileup file was generated using samtools mpileup with parameters *-R -B -q30* and *-Q30*. Using the PileupCaller program from sequenceTools (version 1.5.2) (`https://github.com/stschiff/sequenceTools`), for each SNP, one read covering the SNP was drawn at random, and a pseudohaploid call was made, i.e., the ancient individual was assumed homozygous for the allele on the randomly drawn read for the SNP in question.

##### 1.4.3 Principal component analysis

To carry out the PCA, smartpca (version 16000) (`https:/github.com/DReichLab/EIG`) was used, with the parameters *lsqproject: YES* and *shrinkmode: YES*. Only modern individuals of Eurasian populations and only SNPs on autosomal chromosomes were considered to calculate the principal components. The results of the two individuals that were genotyped in this experiment were only projected onto the calculated components.

##### 1.4.4 Within cluster variance

For each trimming and merging tool and each simulated genome coverage, ten samples were generated and projected on the PCA. To estimate how much the choice of tool influences the accuracy of PCA results at a given coverage, the variance for each cluster of ten samples was calculated using the first ten principal components of the PCA. The within cluster variance, also known as the within-cluster sum of squares, gives an estimation of the dispersion of observations within a cluster, by calculating the average (Euclidean) distance between individual observations and the center of their cluster:

$$\sigma^2 = \frac{1}{n} \sum_{i=1}^{n} \|x_i - \overline{x}\|^2 \qquad (1)$$

where $n$ represents the number of observations, $X$ the total of the observations contained in the cluster and $\overline{x}$ the center of their cluster, also called the centroid.

##### 1.5 Measuring runtime and memory usage

1 million paired-end reads following the insert length distribution of the aDNA library Vi33.19 were simulated and then merged and trimmed with each tool. For AdapterRemoval, bbmerge, fastp and adna-trim, the number of threads was set to one, the other tools did not offer the option to change the thread number. The runtime and memory usage of each tool was reported using snakemake's benchmarking functionality. The measurements were repeated five times and the mean and standard deviation calculated. For the memory usage, the maximum unique set size (USS) is reported, which is the portion of a process's memory that is private to that process and not shared with any other processes. All processes were run on a machine with an Intel Xeon E5620 processor at 2.40GHz and 64402 KB memory.

##### 1.6 Code availability

For each experiment, a workflow was generated using snakemake. Code and workflows used to generate the results of this thesis are available under `https://github.com/liannette/DNA_reconstruct`.

#### 2 Supplementary Results

##### 2.1 Phred accuracy for different quality shifts

To measure the accuracy of the merged Phred quality scores, the theoretical error probabilities of the Phred scores are compared to the observable error rates. Figures 2 to 8 show the Phred quality scores accuracies for the respective $R^2$erging tools at different sequencing qualities (see Section 1.1.3).

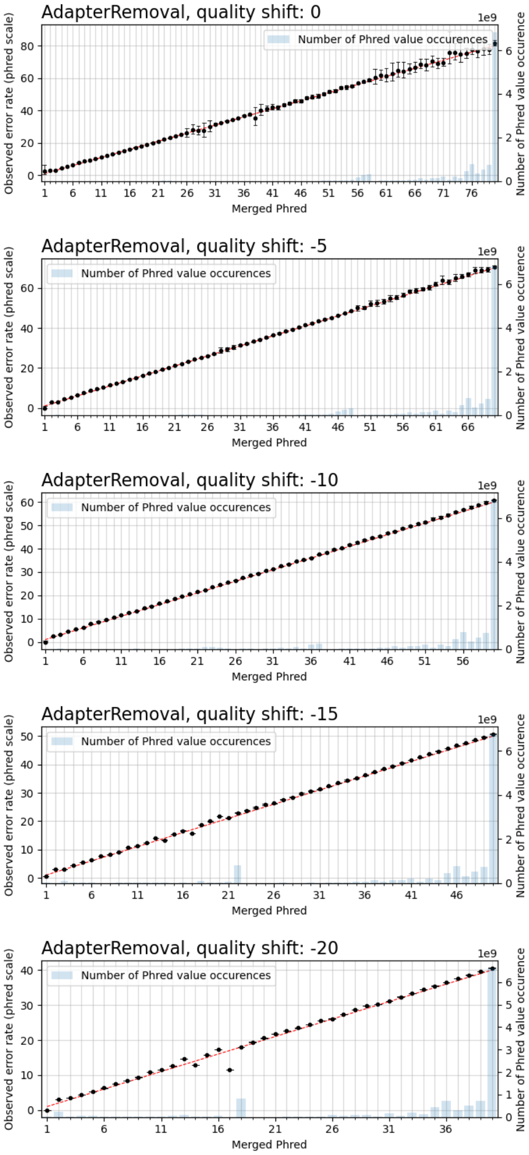


Figure 2: **AdapterRemoval specific observable error rates for merged Phred quality score values at different sequencing qualities.** The Phred-scaled observable error rates are plotted in black. A high Phred value translates to a low error rate. Confidence intervals for the observable error rates were calculated for $\alpha = 0.01$. The red line indicates the perfect fit between observed error rate and merged Phred value, and the number of occurrences of a Phred value in the merged reads is visualized as blue bar plots.

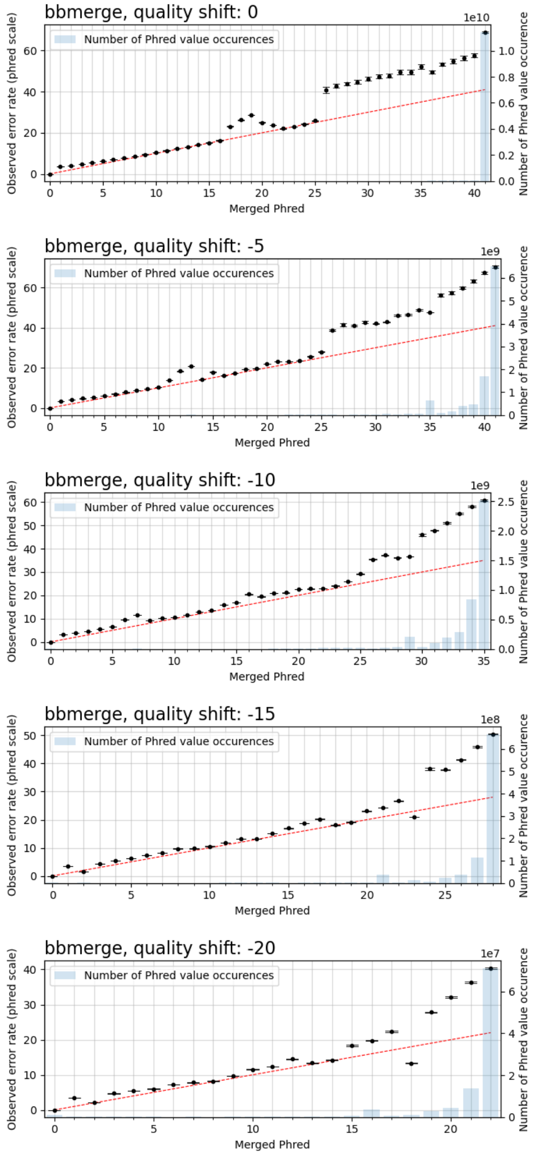


Figure 3: **bbmerge specific observable error rates for merged Phred quality score values at different sequencing qualities.** The Phred-scaled observable error rates are plotted in black. A high Phred value translates to a low error rate. Confidence intervals for the observable error rates were calculated for $\alpha = 0.01$. The red line indicates the perfect fit between observed error rate and merged Phred value, and the number of occurrences of a Phred value in the merged reads is visualized as blue bar plots.

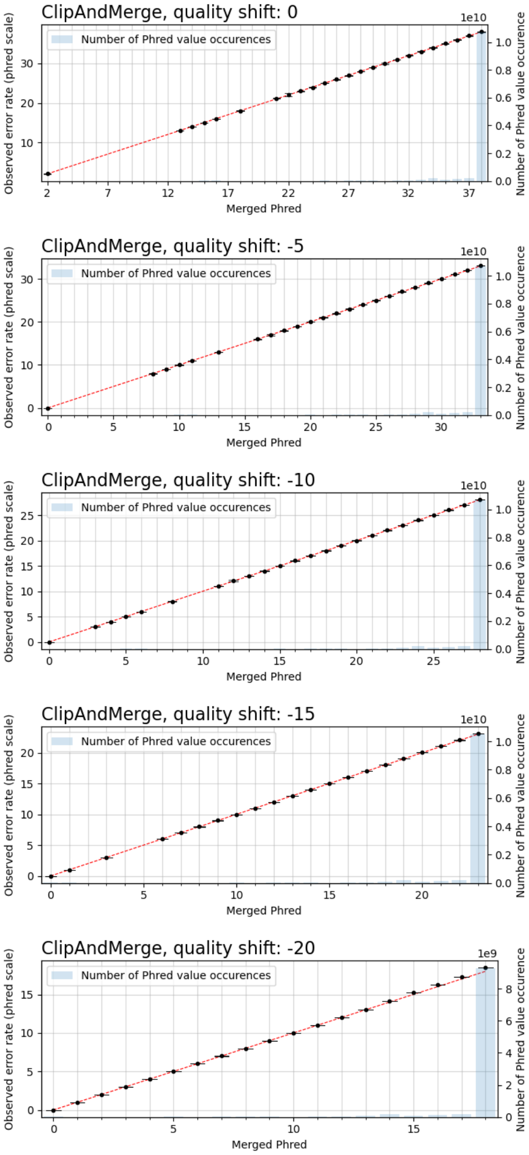


Figure 4: **ClipAndMerge specific observable error rates for merged Phred quality score values at different sequencing qualities.** The Phred-scaled observable error rates are plotted in black. A high Phred value translates to a low error rate. Confidence intervals for the observable error rates were calculated for $\alpha = 0.01$. The red line indicates the perfect fit between observed error rate and merged Phred value, and the number of occurrences of a Phred value in the merged reads is visualized as blue bar plots.

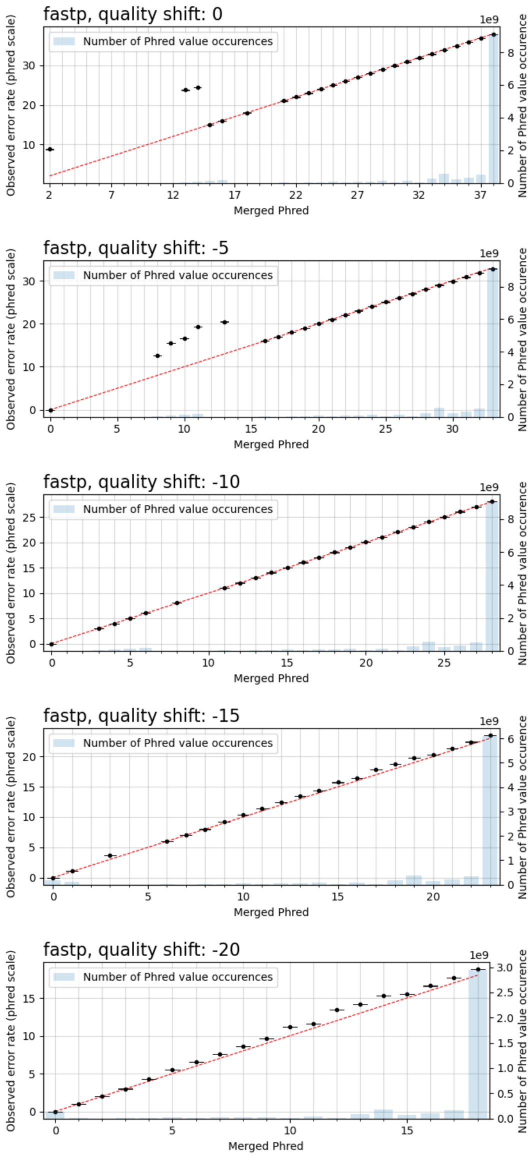


Figure 5: **fastp specific observable error rates for merged Phred quality score values at different sequencing qualities.** The Phred-scaled observable error rates are plotted in black. A high Phred value translates to a low error rate. Confidence intervals for the observable error rates were calculated for $\alpha = 0.01$. The red line indicates the perfect fit between observed error rate and merged Phred value, and the number of occurrences of a Phred value in the merged reads is visualized as blue bar plots.

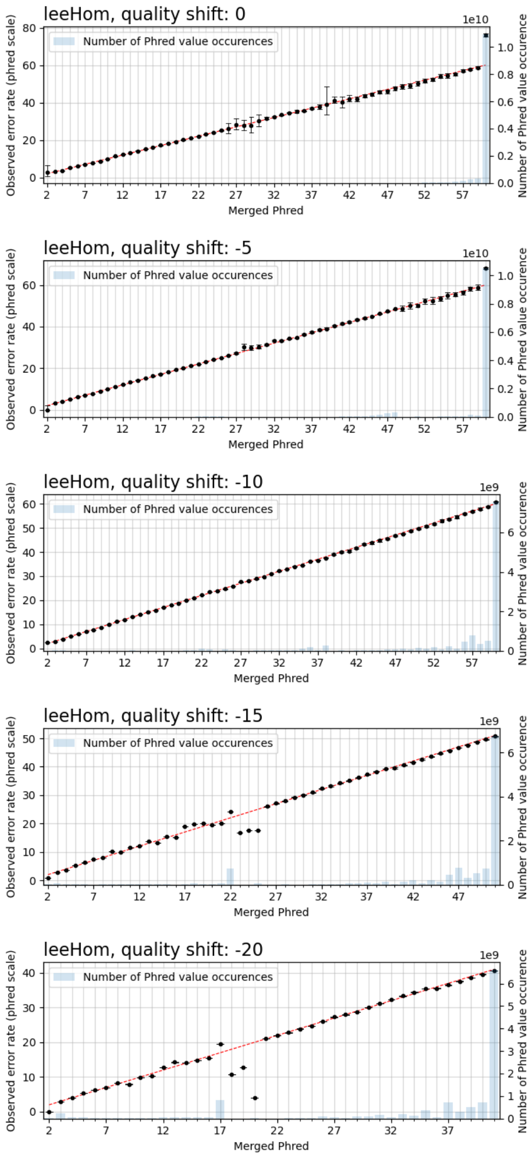


Figure 6: **leeHom specific observable error rates for merged Phred quality score values at different sequencing qualities.** The Phred-scaled observable error rates are plotted in black. A high Phred value translates to a low error rate. Confidence intervals for the observable error rates were calculated for $\alpha = 0.01$. The red line indicates the perfect fit between observed error rate and merged Phred value, and the number of occurrences of a Phred value in the merged reads is visualized as blue bar plots.

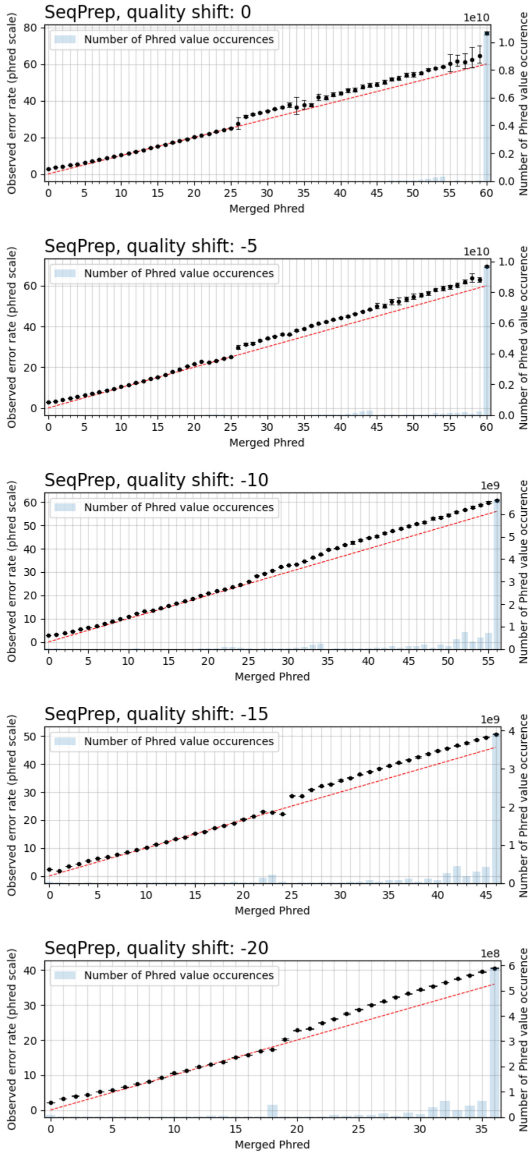


Figure 7: **SeqPrep specific observable error rates for merged Phred quality score values at different sequencing qualities.** The Phred-scaled observable error rates are plotted in black. A high Phred value translates to a low error rate. Confidence intervals for the observable error rates were calculated for $\alpha = 0.01$. The red line indicates the perfect fit between observed error rate and merged Phred value, and the number of occurrences of a Phred value in the merged reads is visualized as blue bar plots.

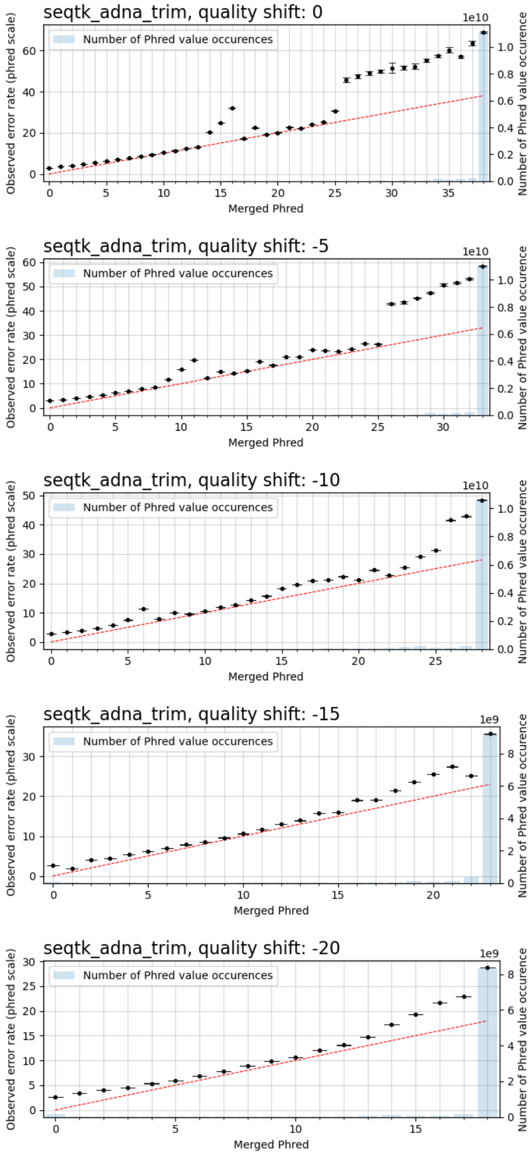


Figure 8: **seqtk/adna_trim specific observable error rates for merged Phred quality score values at different sequencing qualities.** The Phred-scaled observable error rates are plotted in black. A high Phred value translates to a low error rate. Confidence intervals for the observable error rates were calculated for $\alpha = 0.01$. The red line indicates the perfect fit between observed error rate and merged Phred value, and the number of occurrences of a Phred value in the merged reads is visualized as blue bar plots.

#### 2.2 Principal component analysis

The principal components were calculated from genotype data containing 1292 different SNPs from modern day Eurasian populations. The cumulative variance explained by the principal components is shown in figure 9, whereas the individual variance explained by the first 25 principal components is shown in figure 10. To calculate the within cluster variance, the first 10 principal components were used, which cumulative explain 3.12 % of the variance in the dataset. The genotype data for HG002 and HG005, which were generated using the different trimming and merging tool at genome coverages of approximately 0.5X, 1X, 2X and 4X, are projected onto the first two principal components in figure 11 to 14, together with the modern Eurasian populations. At all coverage depths, the projections of all samples for the Ashkenazi Jewish individual (HG002) and the Han Chinese individual (HG005) fall within proximity to their respective populations. In figure 15 to 19, zoomed-in plots are shown for each sample and coverage depth.

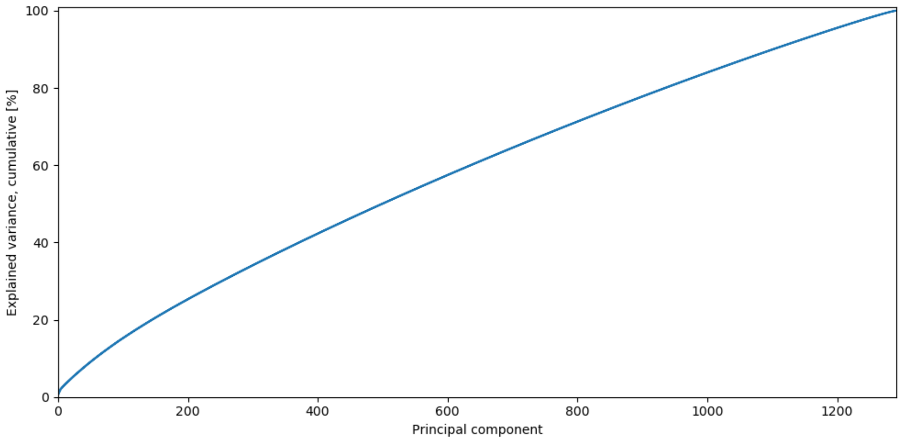


Figure 9: The cumulative variance explained by the principal components.

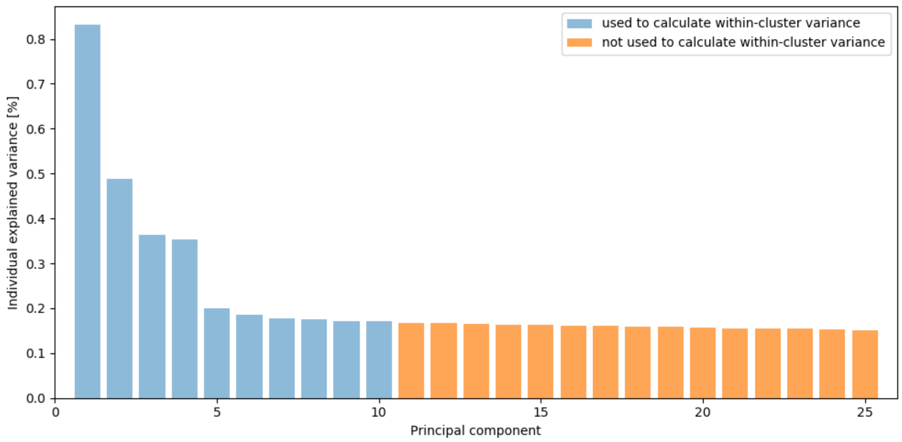


Figure 10: The individual variance explained by the first 25 principal components. The principal components used to calculate the within-cluster variance are plotted in blue.

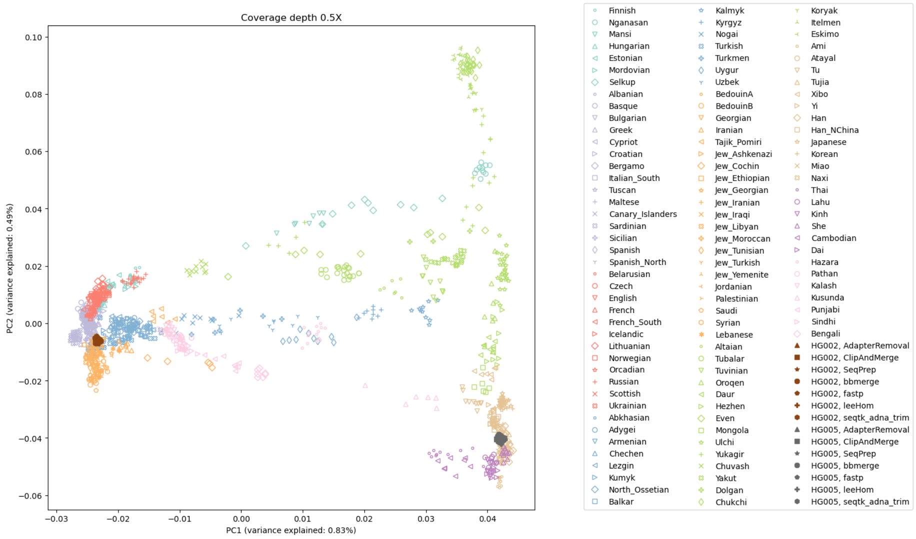


Figure 11: The first two principal components calculated from present-day Eurasian individuals; with projected samples for HG002 and HG005, both at approximately 0.5X reference genome coverage.

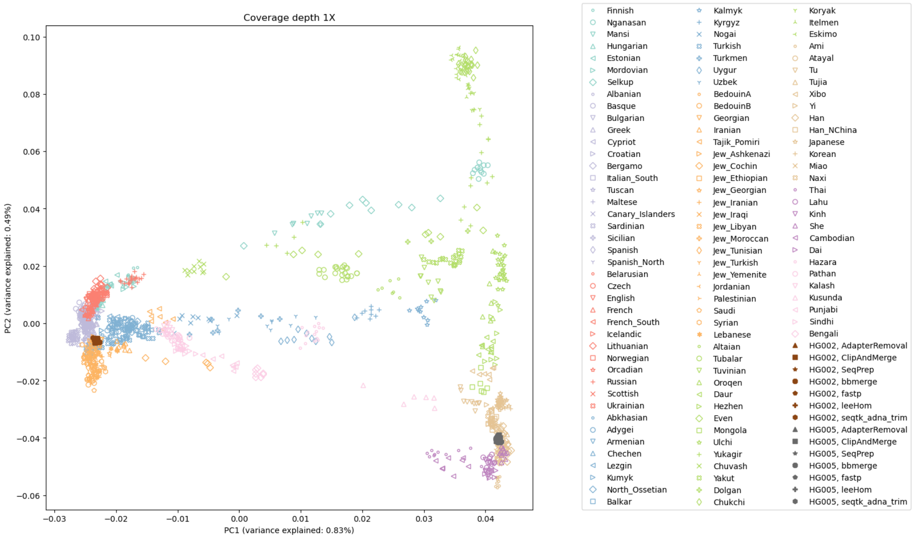


Figure 12: The first two principal components calculated from present-day Eurasian individuals; with projected samples for HG002 and HG005, both at approximately 1X reference genome coverage.

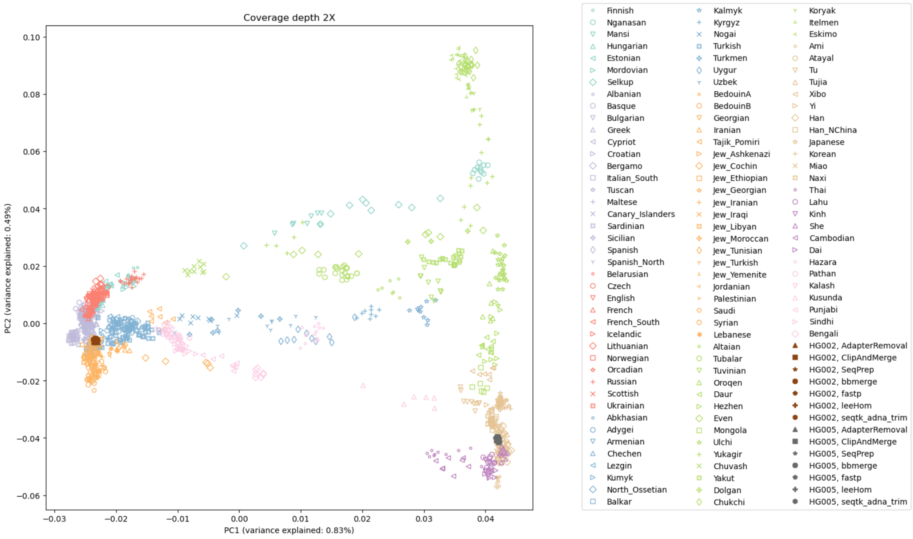


Figure 13: The first two principal components calculated from present-day Eurasian individuals; with projected samples for HG002 and HG005, both at approximately 2X reference genome coverage.

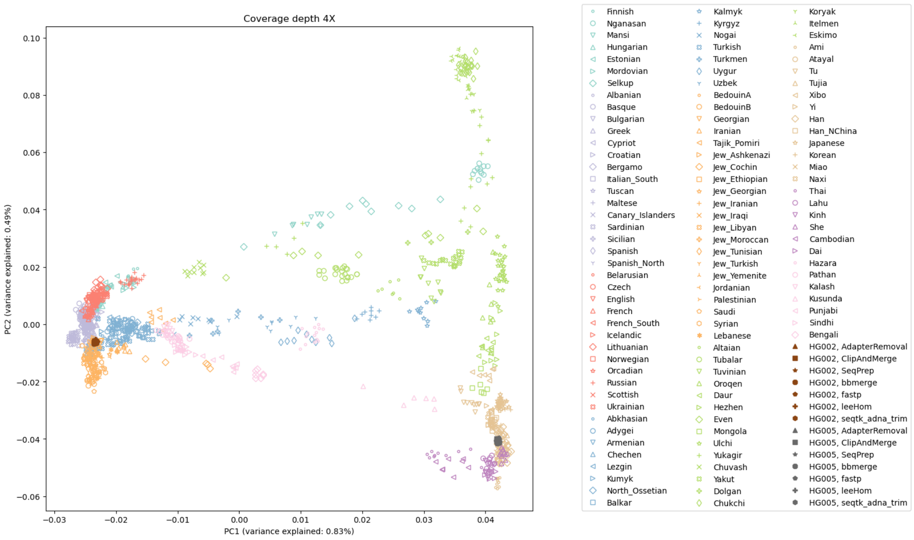


Figure 14: The first two principal components calculated from present-day Eurasian individuals; with projected samples for HG002 and HG005, both at approximately 4X reference genome coverage.

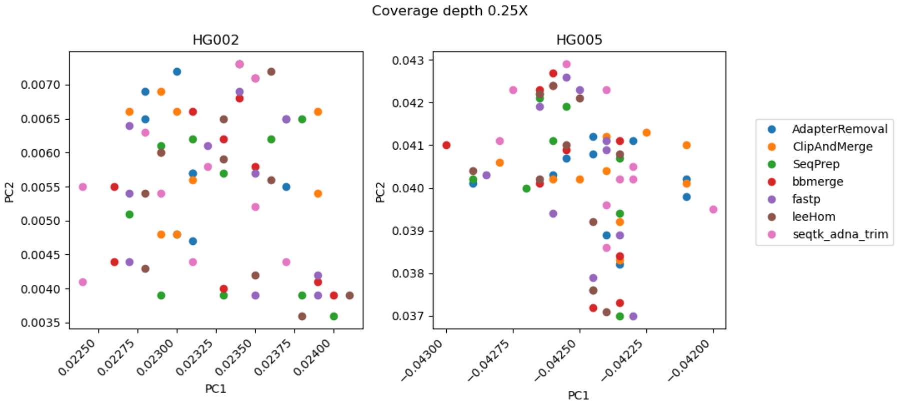


Figure 15: Zoomed-in plots of the first two principal components calculated from present-day Eurasian individuals; with projected samples for HG002 and HG005, both at approximately 0.25X reference genome coverage.

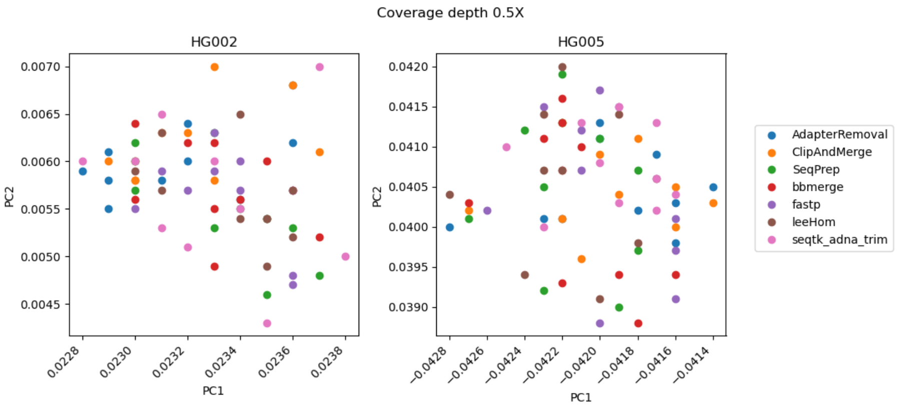


Figure 16: Zoomed-in plots of the first two principal components calculated from present-day Eurasian individuals; with projected samples for HG002 and HG005, both at approximately 0.5X reference genome coverage.

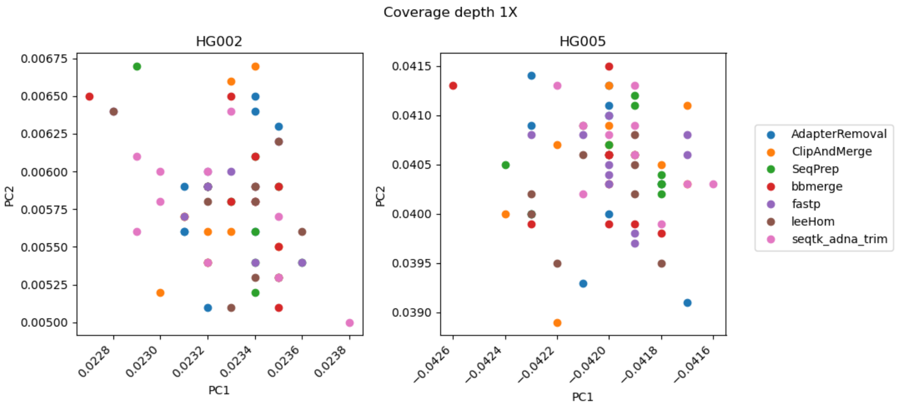


Figure 17: Zoomed-in plots of the first two principal components calculated from present-day Eurasian individuals; with projected samples for HG002 and HG005, both at approximately 1X reference genome coverage.

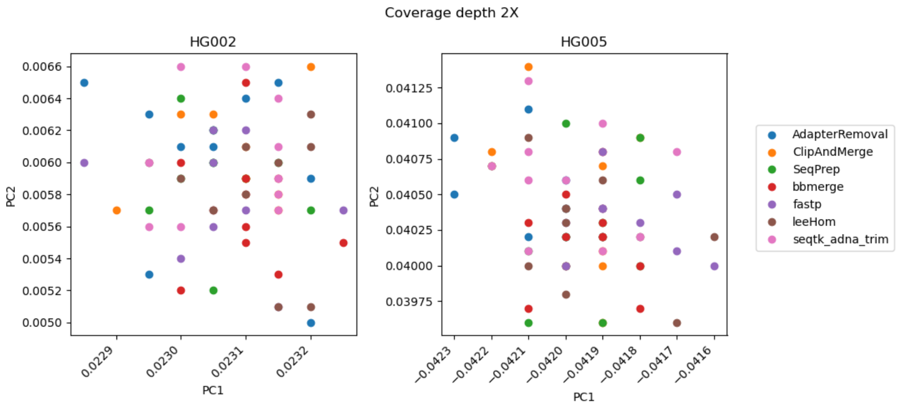


Figure 18: Zoomed-in plots of the first two principal components calculated from present-day Eurasian individuals; with projected samples for HG002 and HG005, both at approximately 2X reference genome coverage.

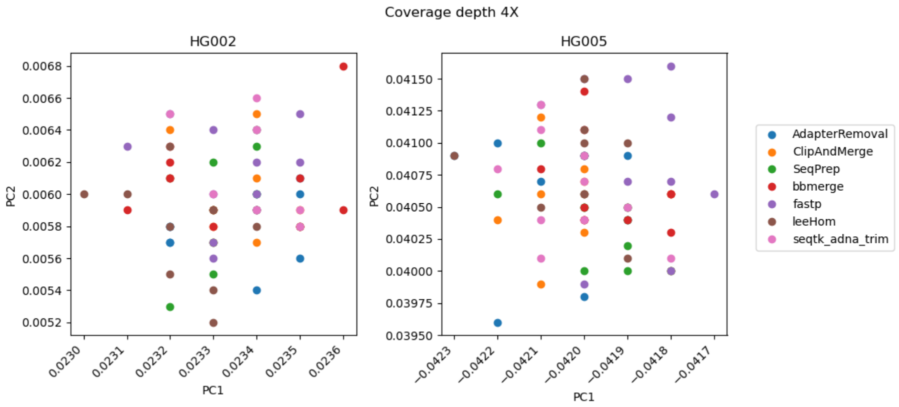


Figure 19: Zoomed-in plots of the first two principal components calculated from present-day Eurasian individuals; with projected samples for HG002 and HG005, both at approximately 4X reference genome coverage.

#### 2.3 Results with non-standardized parameters

The experiments were repeated without specifying the minimum overlap or the minimum length of the merged reads. However, leeHom was still run using the setting for ancient DNA. In figure 20 it can be seen that this affects the range of lengths of the merged reads, with only AdapterRemoval and leeHom producing merged reads shorter than 30 bp. bbmerge has the highest default setting for insert size with 35 bp. The smallest range of merged reads lengths is displayed by fastp with both the minimum insert size and the minimum overlap required for merging and adapter sequence detection being 30 bp. It can be seen in Figure 21 that using the default settings reduced the amount of merged reads when the insert lengths followed the A9180 distribution, as this distribution contains mostly ultrashort sequences, but otherwise it did not visibly affect the merge rate or robustness of the tools for the other insert size distributions. Neither did any tool perform significantly more precise compared to each other using their respective default parameters for minimum length and overlap when performing a PCA for population analysis (figure ).

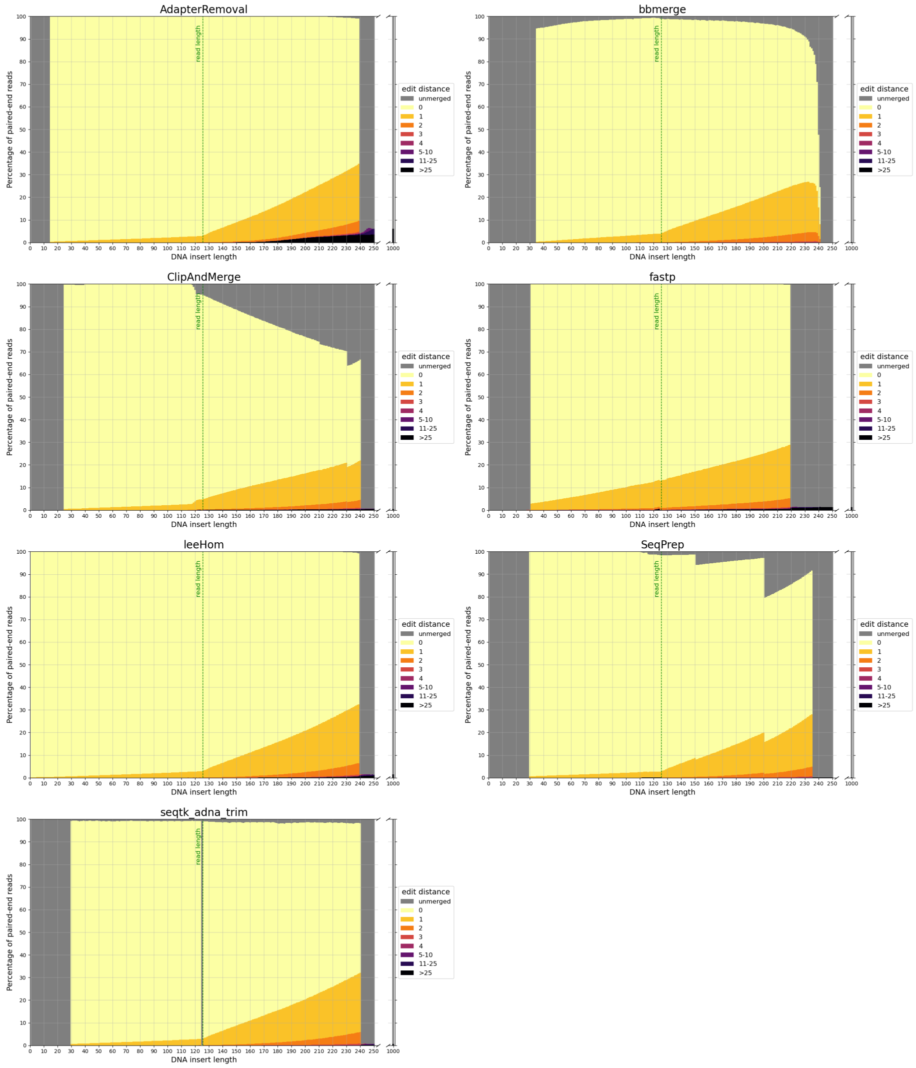


Figure 20: **Sequence accuracy of merged reads for different insert sizes.** Paired-end reads of DNA inserts with different lengths were simulated and then trimmed and merged with the different tools using default parameters for length filtering and minimum overlap length. High edit distances (in darker colors) correspond to high sequence dissimilarity between the merged read and the DNA insert, and unmerged reads are colored in gray. The green line at 125 bp marks the read length. Paired-end reads of inserts with a length up to this value overlap completely. For higher insert lengths, the read overlap decreases, and the proportion of bases that are unique to only one of the paired-end reads increases. For an insert length of 250 bp and higher, the paired-end reads do not overlap.

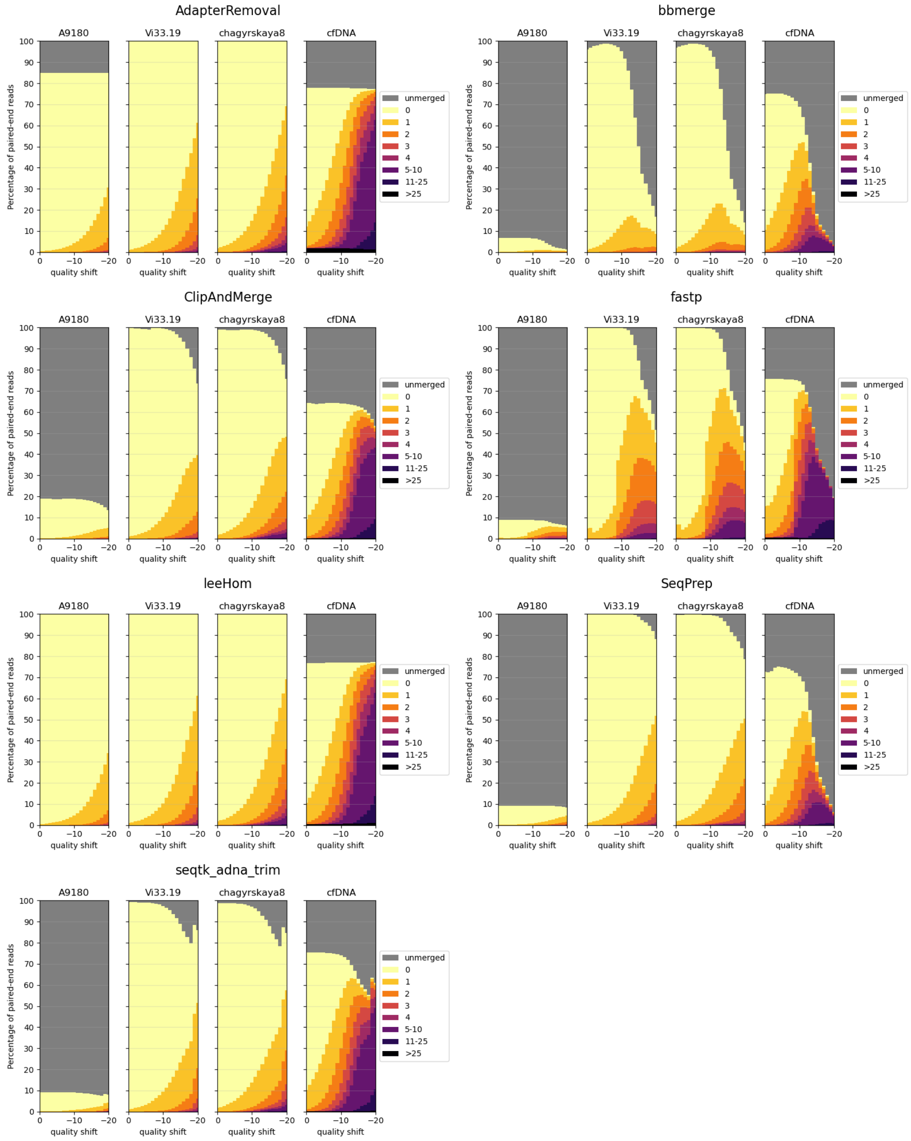


Figure 21: **Sequence accuracy of merged reads for different insert size distributions.** Paired-end reads of DNA inserts with different length distributions were simulated and then trimmed and merged with the different tools using default parameters for length filtering and minimum overlap length. A9189, Vi33.19 and chagyrskaya8 refer to insert size distributions of aDNA, whereas cfDNA represents the insert size distribution of cell-free DNA. High edit distances (in darker colors) correspond to high sequence dissimilarity between the merged read and the DNA insert.

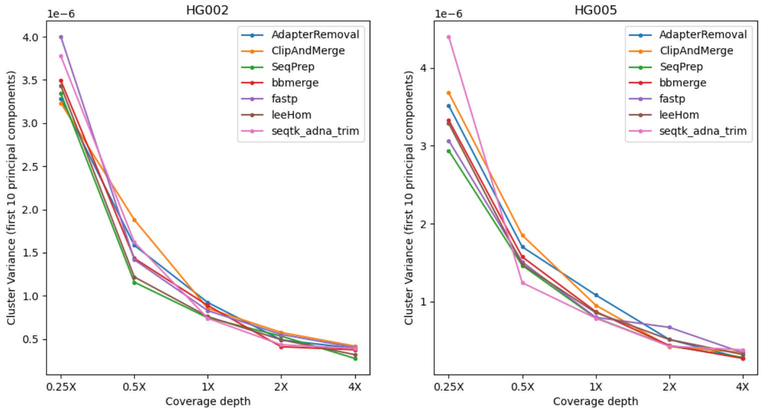


Figure 22: **Within-cluster variance of samples projected onto the PCA.** The trimming and merging tools were used with default parameters for minimum insert size length and minimum overlap length. Each dot represents the within-cluster variance of 10 data points.
